# Dorsal hippocampus contributes to model-based planning

**DOI:** 10.1101/096594

**Authors:** Kevin J. Miller, Matthew M. Botvinick, Carlos D. Brody

## Abstract

Planning can be defined as a process of action selection that leverages an internal model of the environment. Such models provide information about the likely outcomes that will follow each selected action, and their use is a key function underlying complex adaptive behavior. However, the neural mechanisms supporting this ability remain poorly understood. In the present work, we adapt for rodents recent advances from work on human planning, presenting for the first time a task for animals which produces many trials of planned behavior per session, allowing the experimental toolkit available for use in trial-by-trial tasks for rodents to be applied to the study of planning. We take advantage of one part of this toolkit to address a perennially controversial issue in planning research: the role of the dorsal hippocampus. Although prospective representations in the hippocampus have been proposed to support model-based planning, intact planning in hippocampally damaged animals has been observed in a number of assays. Combining formal algorithmic behavioral analysis with muscimol inactivation, we provide the first causal evidence directly linking dorsal hippocampus with planning behavior. The results reported, and the methods introduced, open the door to new and more detailed investigations of the neural mechanisms of planning, in the hippocampus and throughout the brain.

## Introduction

Imagine a game of chess. As the players think about their next moves, they consider the outcome each action would have on the board, as well as the opponent’s likely reply. The players’ knowledge of the board and the rules constitutes an *internal model* of chess, a knowledge structure that links actions to their likely outcomes. The process of using such an “action-outcome” model to inform behavior is defined within reinforcement learning theory as the act of *planning*^1^. We will follow this definition here. Planning, so defined, has been an object of scientific investigation for many decades and in many subfields, and the resulting studies have generated important insights into the planning abilities of both humans and other animals^2–5^. However, despite such progress, the neural mechanisms that underlie planning remain frustratingly obscure.

One important reason for this continuing uncertainty lies in the behavioral assays that have traditionally been employed. Until recently, our understanding of the neural mechanisms of planning^2,6–8^ has been informed largely by behavioral tests (such as outcome devaluation) in which the subject is put through a sequence of training stages and then makes just one single decision to demonstrate planning (or an absence thereof). The same animal can be tested multiple times^9^, but at most one behavioral measure of planning is obtained per session. Seminal studies using these assays have established the relevance of several neural structures^3,4^, and they continue to be fundamental for many experimental purposes. But these assays are constrained by the fact that they elicit only a small number of planned decisions per experiment. In an important recent breakthrough, new tasks have been developed that lift this constraint^10–13^, allowing the collection of many repeated trials of planned behavior. The new tasks provide an important complement to existing behavioral assays, promising to allow both a detailed evaluation of competing models as well as new opportunities for experiments investigating the neural mechanisms of planning. However, these tasks have so far been applied only to human subjects, limiting the range of experimental techniques which can be employed.

In the present work, we adapted for rats a multi-trial decision task from a recent influential line of research in humans^10^. Using behavioral data from a first experiment, we conducted a set of detailed computational analyses, in order to confirm that rats, like humans, employ model-based planning to solve the task. In a second experiment, taking advantage of the opportunity to employ causal neural techniques not available in humans, we used the task to address an important open question in the neuroscience of planning: the role of the dorsal hippocampus.

A long-standing theory of hippocampal function holds that it represents a “cognitive map” of physical space used in support of navigational decision-making^14^. Classic experiments demonstrate behavioral impairments on navigation tasks resulting from hippocampal damage^15,16^, as well as the existence of “place cells” which both encode locations in space^17^ and “sweep out” potential future paths at multiple timescales-. These findings have given rise to computational accounts of hippocampal function that posit a key role for the region in model-based planning^22–25^. However, support for these theories from experiments employing causal manipulations has been equivocal. Studies of both spatial navigation and instrumental conditioning have shown intact action-outcome behaviors following hippocampal damage^26–31^. At the same time, tasks requiring relational memory do show intriguing impairments following hippocampal damage^32–35^. The latter tasks assay whether behavior is guided by knowledge of relationships between stimuli (stimulus-stimulus associations), which plausibly involve similar representations and structures as the action-outcome associations that underlie planning, but they do not focus on action-outcome associations specifically. Here, with the two-step task, our focus is on these latter types of associations.

Using rats performing the two-step task, we performed reversible inactivation experiments in both dorsal hippocampus (dH) and in orbitofrontal cortex (OFC), a brain region widely implicated in model-based control in traditional assays^36–39^. The repeated-trials nature of the task allows us to use computational modeling to identify a set of separable behavioral patterns (such as model-based planning versus model-free control) which jointly explain observed behavior, and to quantify the relative strength of each pattern. We find that the behavior of our animals is dominated by a pattern consistent with model-based planning, with important influences of simpler patterns including novelty aversion, perseveration, and bias. The model-based pattern is selectively impaired by inactivation of OFC or dH, while other behavioral patterns are unaffected.

Importantly, model-based planning depends on a number of computations – behaviorally observed planning impairments might be caused by impairments to the planning process itself or instead by impairments to learning and memory processes upon which planning depends. Computational modeling analysis indicates that our effects are not well-described as an impairment in learning or memory in general, but as a specific attenuation of model-based behavior. We therefore conclude that these regions either perform computations integral to the planning process itself (i.e. use of the action-outcome model to inform choice) or represent inputs that are used specifically by the planning process. This provides what is, to our knowledge, the first causal evidence that disabling the dorsal hippocampus impairs model-based planning behavior.

## Results

We trained rats to perform a multi-trial decision making task^10^ adapted from the human literature on model-based control (Fig. 1), where it is widely referred to as the ‘two-step task.’ The two-step task is designed to distinguish model-based versus model-free behavioral strategies. In the first step of this task, the rat chooses between two available choice ports, each of which will lead to one of two reward ports becoming available with probability 80% (*common transition*), and to the other reward port with probability 20% (*uncommon transition*). In the second step of the task, the rat does not have a choice, but is instead instructed as to which reward port has become available, enters it, and either receives or does not receive a water reward. Reward ports differ in the probability with which they deliver reward, and reward probability changes at unpredictable intervals (see Methods). Optimal performance requires learning which reward port currently has the higher reward probability, and selecting the choice port more likely to lead to that port. This requires using knowledge of the likely outcomes that follow each possible chosen action – that is, it requires planning.

Rats performed the two-step task in a behavioral chamber outfitted with six nose ports arranged in two rows of three (Fig. 1). Choice ports were the left and right side ports in the top row, and reward ports were the left and right side ports in the bottom row. Rats initiated each trial by entering the center port on the top row, and then indicated their choice by entering one of the choice ports. This resulted in an auditory stimulus playing, which indicated which of the two reward ports was about to become available. Before the reward port became available, however, the rat was required to enter the center port on the bottom row. This was done in order to keep motor acts relatively uniform across common and uncommon trial types. For some animals, the common transition from each choice port led to the reward port on the same side (as in Figure 1A; *common-congruent* condition), while for others, it led to the reward port on the opposite side (*common-incongruent*). The transition probabilities for a particular animal were kept fixed throughout that animal’s entire experience with the task. These probabilities are therefore likely to be stable features of internal mappings from actions to their likely outcomes, i.e., stable features of subjects’ internal models of the world.

We trained 21 rats to perform the two-step task in daily behavioral sessions (n=1959 total sessions), using a semi-automated training pipeline which enabled us to run large numbers of animals in parallel with minimal human intervention (see Methods, *Behavioral Training Pipeline*). This formalization of our training procedure into a software pipeline should also facilitate efforts to replicate our task in other labs, since the pipeline can readily be downloaded and identically re-run.

**Figure 1:**
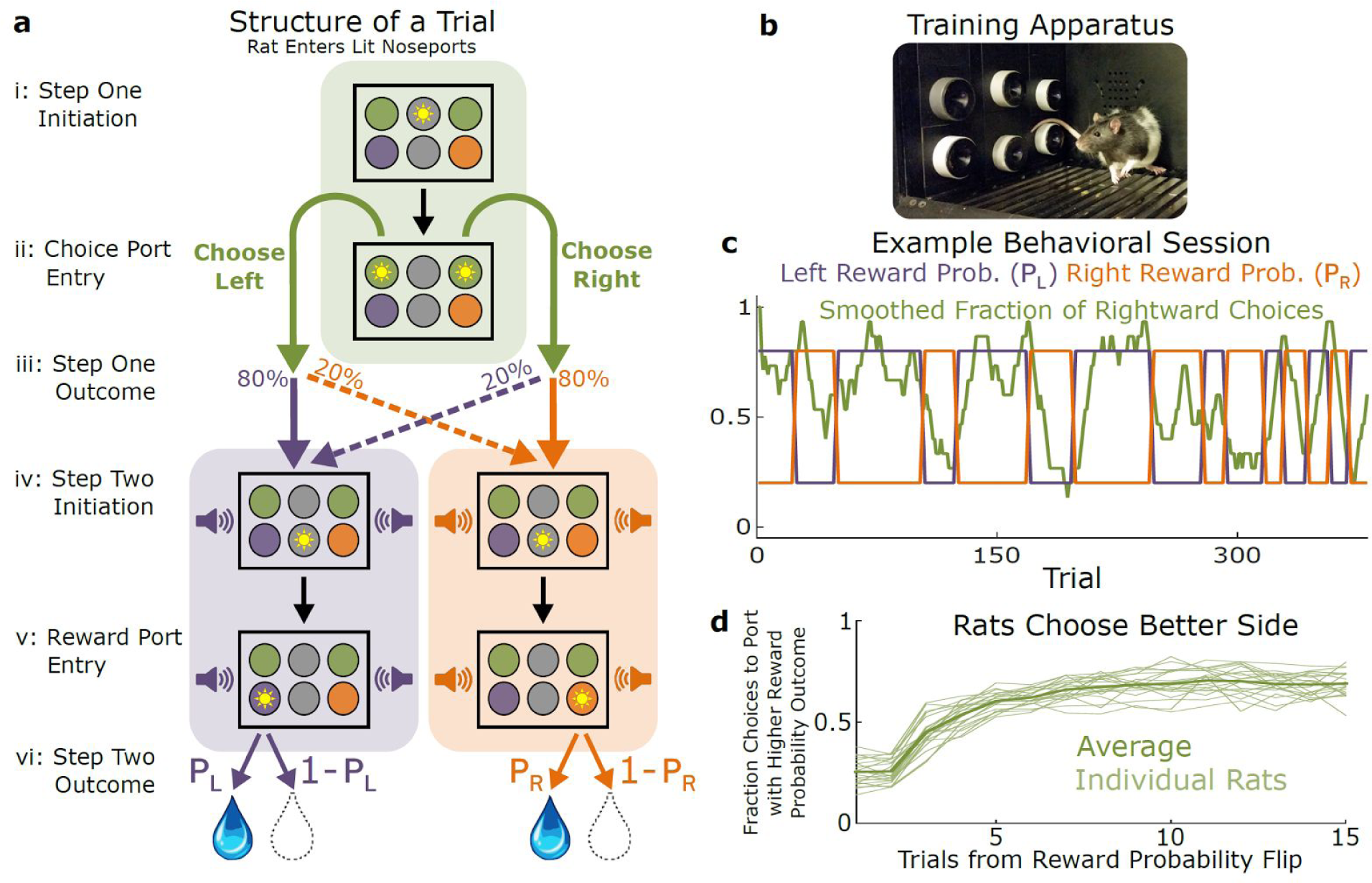
Two-Step DecisionTask for Rats. A) Structure of a single trial of the two-step task. i) Top center port illuminates to indicate trial is ready, rat enters it to initiate the trial. ii) Choice ports illuminate, rat indicates decision by entering one of them. iii) Probabilistic transition takes place, with probability depending on the choice of the rat. Sound begins to play, indicating the outcome of the transition. iv) Center port in the bottom row illuminates, rat enters it. v) The appropriate reward port illuminates, rat enters it. vi) Reward is delivered with the appropriate probability. B) Photograph of behavioral apparatus, consisting of six nose-ports with LEDs and infrared beams, as well as a speaker mounted in the rear wall. C) Example behavioral session. Rightward choices are smoothed with a 10-trial boxcar filter. At unpredictable intervals, reward probabilities at the two ports flip synchronously between high and low. Rats adapt their choice behavior accordingly. D) Choice data for all rats. The fraction of trials on which the rat selected the choice port whose common (80%) transition led to the reward port with currently higher reward probability, as a function of the number of trials that have elapsed since the last reward probability flip.

## Two analysis methods to characterize behavior and quantify planning

Although optimal performance in the two-step task requires a model-based strategy, good performance can be achieved by both model-based and model-free strategies, with similar reward rates (Supp Fig 1). Critically, however, these two types of strategies can be distinguished by the patterns of choices made by the subject^10^. Model-free strategies tend to repeat choices that ultimately resulted in reward, and to avoid choices that led to reward omission, regardless of whether the transition after the choice was a common or an uncommon one. This is because model-free strategies, by definition, do not use knowledge of action-outcome probabilities. In contrast, model-based planning strategies, which do use knowledge of the causal structure of the environment, are sensitive to whether the transition following a choice was a common or an uncommon one. Thus, after an uncommon transition, model-based strategies tend to avoid choices that led to a reward, because the best way to reach the rewarding port again is through the common transition that follows the opposite choice. Similarly, again after an uncommon transition, model-based strategies tend to repeat choices that led to a reward omission, because the best way to avoid the unrewarding port is through the common transition likely to occur after repeating the choice. Both of these patterns are in contrast to model-free strategies. Following this logic, Daw et al. (2011) examined how humans’ choices in a given trial depend on the immediately previous trial, and concluded that humans appear to use a mixture of model-based and model-free strategies^10^.

To assess the extent to which rat subjects were using a model-based strategy, we extended the analysis of Daw et al. (2011), which considered the influence of the immediately preceding trial on present-trial behavior, to now use information from multiple trials in the past. We have shown separately that that this many-trials-back approach is robust to some potential artifacts that are an issue for the one-trial-back approach (one of which, for example, would be due to slow learning rates; cite Bar Plots Paper). The many-trials-back approach consists of a logistic regression model that predicts the probability with which a rat will select the left or the right choice port on a particular trial, given the history of recent trials. A trial that occurred τ trials ago can be one of four types: common-rewarded, uncommon-rewarded, common-omission, and uncommon-omission. For each *τ*, each of these trial types is assigned a weight (*β*_*CR*_(τ), *β*_*UR*_(τ), *β*_*CO*_(τ), *β*_*UO*_(τ) respectively). Positive weights correspond to greater likelihood to make the same choice that was made on a trial of that type which happened *τ* trials in the past, while negative weights correspond to greater likelihood to make the other choice. The weighted sum of past trials’ influence then dictates choice probabilities (see Methods, *Behavior Analysis*, equation one). This model contains one hyperparameter, *T*, giving the number of past trials τ that will be considered. Larger values of *T* allow the model to consider longer time dependence in behavior, at the expense of adding additional model complexity. We found that for our rats, increasing *T* beyond 5 resulted in negligible increases in quality of fit (Figure S2). For each subject, we used maximum likelihood fitting to find the model weights that best matched that subject’s choices. The resulting model weights quantify the subject’s tendency to choose the same choice port that it has selected in the past vs. choose the other port, as a function of the number of intervening trials, the choice made (left vs. right port), the first-step outcome (common vs. uncommon transition), and the second-step outcome (reward vs. omission). Importantly, because model-free strategies do not distinguish between common and uncommon transitions, model-free strategies will tend to have *β*_CR_ ≈ *β*_UR_ and *β*_CO_ ≈ *β*_UO_. In contrast, model-based strategies tend to change their behavior in different ways following common versus uncommon transitions, and will therefore have *β*_CR_ > *β*_UR_ and *β*_CO_ < *β*_UO_.

Applying this approach to synthetic data from artificial reinforcement learning agents using planning-based or model-free strategies (see Methods: *Synthetic Behavioral Data*) yields the expected patterns (Fig. 2A,B). For the planning agent (Fig. 2A), trials with common (solid) and uncommon (dashed) transitions have opposite effects on the current choice (compare blue solid versus blue dashed lines, and compare red solid versus red dashed). In contrast, for the model-free agent (Fig. 2B), common and uncommon transitions have the same effect (solid and dashed lines overlap), and only reward versus reward omission is important (red versus blue curves). Figure 2C shows the result of fitting the regression model to data from an example rat. The behavioral patterns observed are quite similar to those expected of a model-based agent (compare 2A to 2C).

**Figure 2:**
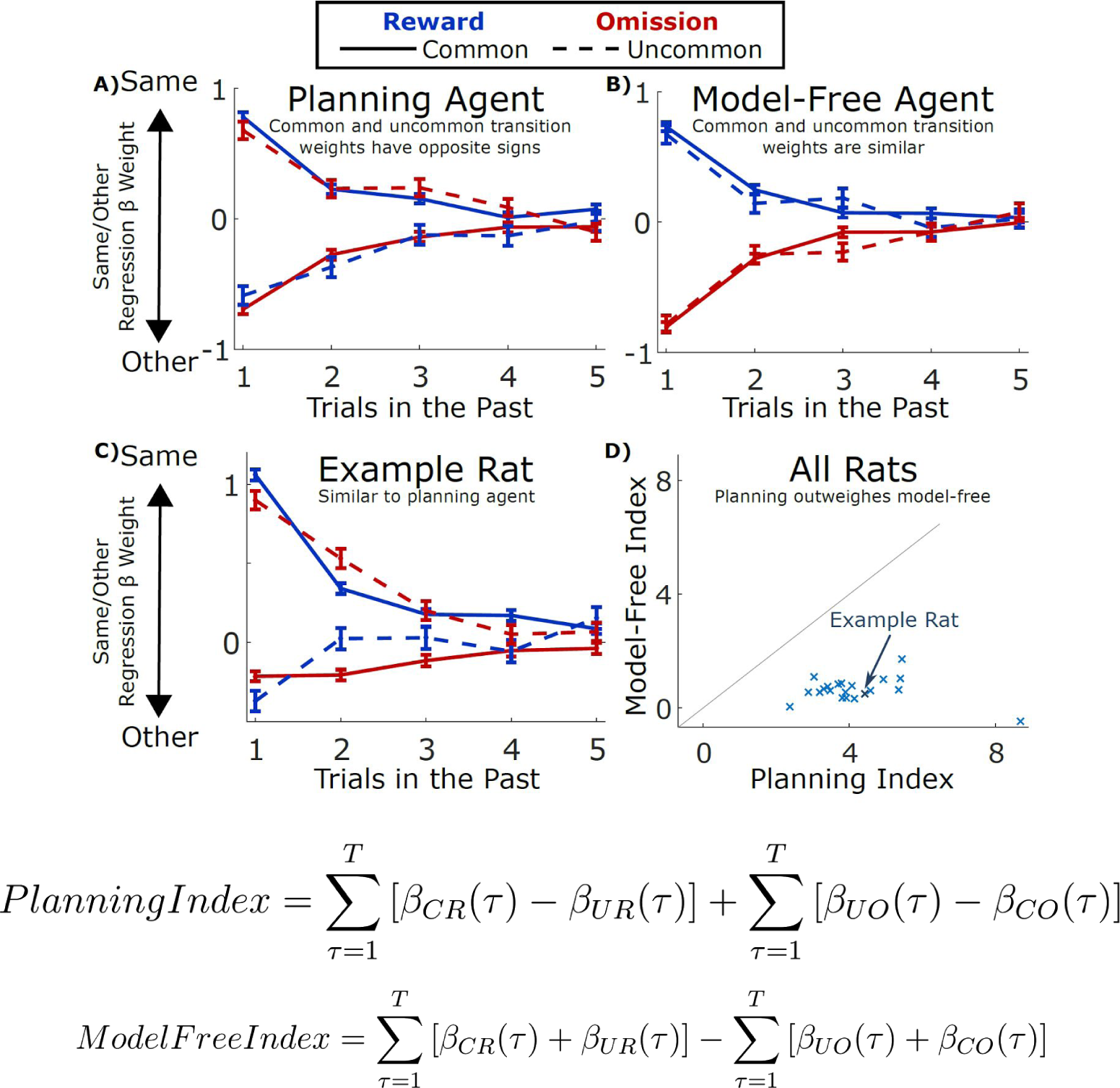
Behavior Analysis Overview. A) Results of the trial-history regression analysis applied to simulated data from a model-based planning agent, and from (B) a model-free temporal difference learning agent. C) Results of the analysis applied to data from an example rat. D) Model-free and planning indices computed from the results of the regression analysis, shown for all rats in the dataset.

We applied the above regression analysis to the behavior of each rat in our dataset (Figure S3) to reveal the nature of each animal’s choice strategy. To quantify the overall extent to which each rat’s dataset showed evidence of a planning vs. a model-free strategy, we defined a “planning index” and a “model-free index” by summing over the regression weights consistent with each pattern (see Fig. 2; Methods, *Behavior Analysis*). We have found that these measures provide a more reliable guide to behavioral strategy than standard measures, which consider only the immediately previous trial (see Miller, Brody, & Botvinick, 2016, *bioRxiv*, for details^40^). We found that trained rats overwhelmingly showed large positive planning indices (see Figure 2; mean over rats: 4.2, standard error 0.3), and small positive model-free indices (mean: 0.6, standard error 0.1), strongly suggesting that they had adopted a planning strategy. We also found that movement times from the bottom center port to the reward port were faster for common vs. uncommon transition trials (average median movement time 700 ms for common and 820 ms for uncommon, p < 10^−5^; Figure S3), further supporting the idea that rats used knowledge of the transition probabilities to inform their behavior. These results were similar between rats in the common-congruent condition (common outcome for each choice port is the reward port on the same side, as in Figure 1A) and those in the common-incongruent condition (common outcome is the port on the opposite side; p>0.2).

This regression analysis also revealed first, that there is substantial rat-by-rat variability (Figure 3A, top panel), and second, that there are important deviations from the predicted model-based pattern (Figure 3A; Figure S4). For example, the rat in the top left panel of Fig. 3A (same rat as in Fig. 2C) shows the overall pattern of regression weights expected for a model-based strategy, but in addition all weights are shifted slightly in the positive direction (i.e. the “repeat choice” direction). This particular rat’s behavior can thus be succinctly described as a combination of a model-based strategy plus a tendency to repeat choices; we refer to the latter behavioral component as “perseveration”. Many other behavior components are also possible, including win-stay/lose-switch, response bias, and others. While the regression analysis’ rich and relatively theory-neutral description of each rat’s behavioral patterns is useful for identifying such deviations from a purely model-based strategy, it is limited in its ability to disentangle the extent to which each individual deviation is present in a dataset. The regression analysis suffers from several other disadvantages as well – it requires a relatively large number of parameters (21 weights for a model with T=5), and it is implausible as a generative account of the computations used by the rats to carry out the behavior (requiring an exact memory of the past five trials). We therefore turned to a complementary analytic approach: trial-by-trial model fitting using mixture-of-agents models.

Mixture-of-agents models provide both more parsimonious descriptions of each rat’s dataset (involving fewer parameters) and more plausible hypotheses about the underlying generative mechanism. Each model comprises a set of agents, each deploying a different choice strategy. Rats’ observed choices are modeled as reflecting the weighted influence of these agents, and fitting the model to the data means setting these weights, along with other parameters internal to the agents, so as to best match the observed behavior. We limited the memory associated with each agent to only a single scalar variable (in contrast to the exact five-trial memory of the regression model) greatly increasing the biological plausibility of each agent. This also makes the mixture models far less flexible than the full regression model. Despite this, we nevertheless found that a mixture-of-agents model could account very well for the rats’ behavior (compare top row to bottom row, Fig. 3A). We found that a good qualitative match to rats’ behavior could be achieved with a mixture of only four simple agents, representing four patterns – we call these patterns planning, choice perseveration, novelty preference, and choice bias (Figure 3A; Figure S4). The four agents implementing these four patterns were a model-based reinforcement learning agent (planning), an agent which repeated the previous trial’s choice (perseveration), an agent which repeated or avoided choices which led to a common vs. an uncommon transition (novelty preference), and an agent which prefered the left or the right choice port on each trial (choice bias; see Methods, *Behavioral Models*). In all, this model contained five free parameters: four mixing weights, *β*_plan_, *β*_np_, *β*_persev_, and *β*_bias_, associated with each of the four agents, as well as a learning rate, *α*_plan_, internal to the planning agent. This mixture-of-agents model was fit to individual rats’ datasets using maximum *a posteriori* fitting under weakly informative priors (see Methods, *Behavioral Model Fitting*). We arrived at these four particular patterns as the necessary components because removing any of the four agents from the mixture resulted in a large decrease in quality of fit (assessed by cross-validated likelihood: Fig 3B, red; Methods, *Model Comparison*), and because adding a variety of other additional patterns (model-free reinforcement learning, model-based and model-free win-stay/lose-switch, transition learning, or all of the above; Methods, *Behavioral Models*) resulted in only negligible improvements (Fig 3B, green). Substituting an alternate learning mechanism based on Hidden Markov Models into the planning agent also resulted in a negligible change (Fig3B, blue; Methods, *Behavioral Models*), and we do not consider the HMM learner further. We found that the mixture model performed similarly in quality of fit to the regression-based model, for all but a minority of rats (Fig 3B, blue). We computed normalized mixing weights for each agent for each rat, quantifying the extent to which the behavior of that rat was driven by the influence of that agent, with all other agents taken into account (Fig 3C). The planning agent earned the largest mixing weight for each rat, indicating that model-based planning is the dominant component of behavior on our task. Taken together, these findings indicate that this mixture model is an effective tool for quantifying patterns present in our behavioral data, and that well-trained rats on the two-step task adopt a strategy dominated by model-based planning. However, rats also exhibit perseveration, novelty preference, and bias.

**Figure 3:**
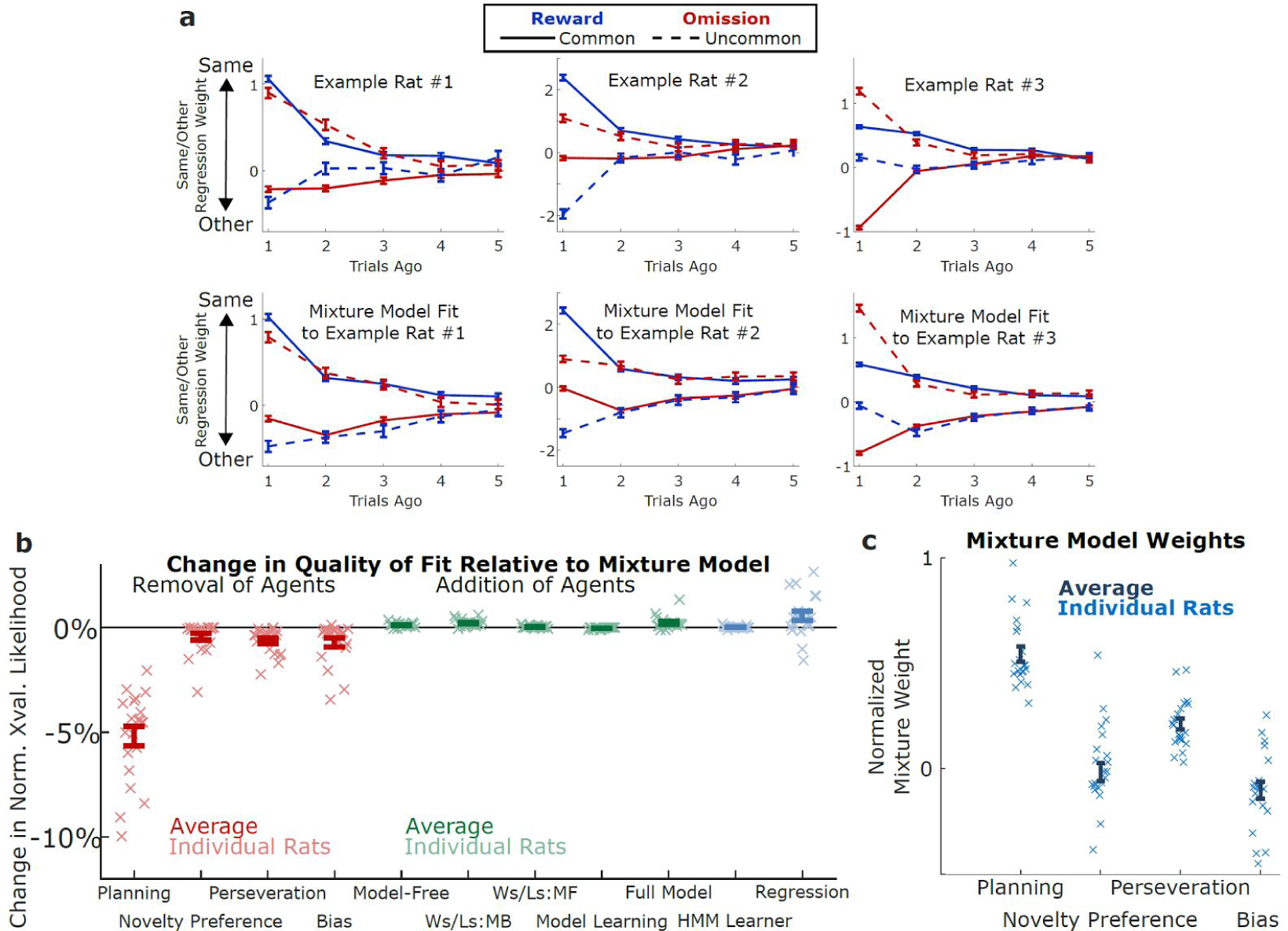
Model-Fitting Analysis. A) Results of the trial-history regression analysis applied to data from three example rats (above) and simulated data produced by the agent model with parameters fit to the rats (below) B) Change in quality of fit resulting from removing (red) or adding (green) components to the reduced behavioral model. C) Normalized mixture weights resulting from fitting the model to rats’ behavior

## Pharmacological inactivation of hippocampus or orbitofrontal cortex impairs planning

In the next phase of this work, we took advantage of the regression analysis and the mixture-of-agents model as computational tools for examining the causal contribution of OFC and dH on particular components of behavior. We implanted six well-trained rats with infusion cannulae bilaterally targeting each region, and used these cannulae to perform reversible inactivations, infusing the GABA-A agonist muscimol (a pharmacological agent that silences neural activity for a period of many hours^41,42^), then allowing the animals to recover for a short time before placing them in the behavioral chamber to perform the task (see Methods, *Inactivation Experiments*). We compared behavior during muscimol sessions to behavior during control sessions performed the day before and after inactivation, as well as to sessions in which we infused saline into the target regions (see Figures S5 and S6). We computed the regression-based planning index for each rat (as in Fig. 2), and found that inactivation of either region substantially reduced the magnitude of the planning index relative to each region’s control sessions (Figure 4; OFC p=0.001, dH p=0.01, see Methods, *Analysis of Inactivation Data*), as well as to pooled saline sessions (OFC p=0.004, dH, p=0.04). We found no effect of either inactivation on the model-free index (all p > 0.5). The impact of inactivation on model-based behavioral patterns was not simply due to an overall reduction of the modulation of past trials on current trial choices: we computed the aggregate main effect of past choices on future choices (*β*_CR_ + *β*_UR_ + *β*_CO_ + *β*_UO_) for each rat for each type of session, and found that this feature of the data was insensitive to inactivation of either region (Figure 4b; OFC p=0.4, dH p=0.7). Together, these results suggest that inactivation of OFC or dH reduces the extent to which behavior shows evidence of planning, but does not affect evidence for perseveration or model-free patterns.

**Figure 4:**
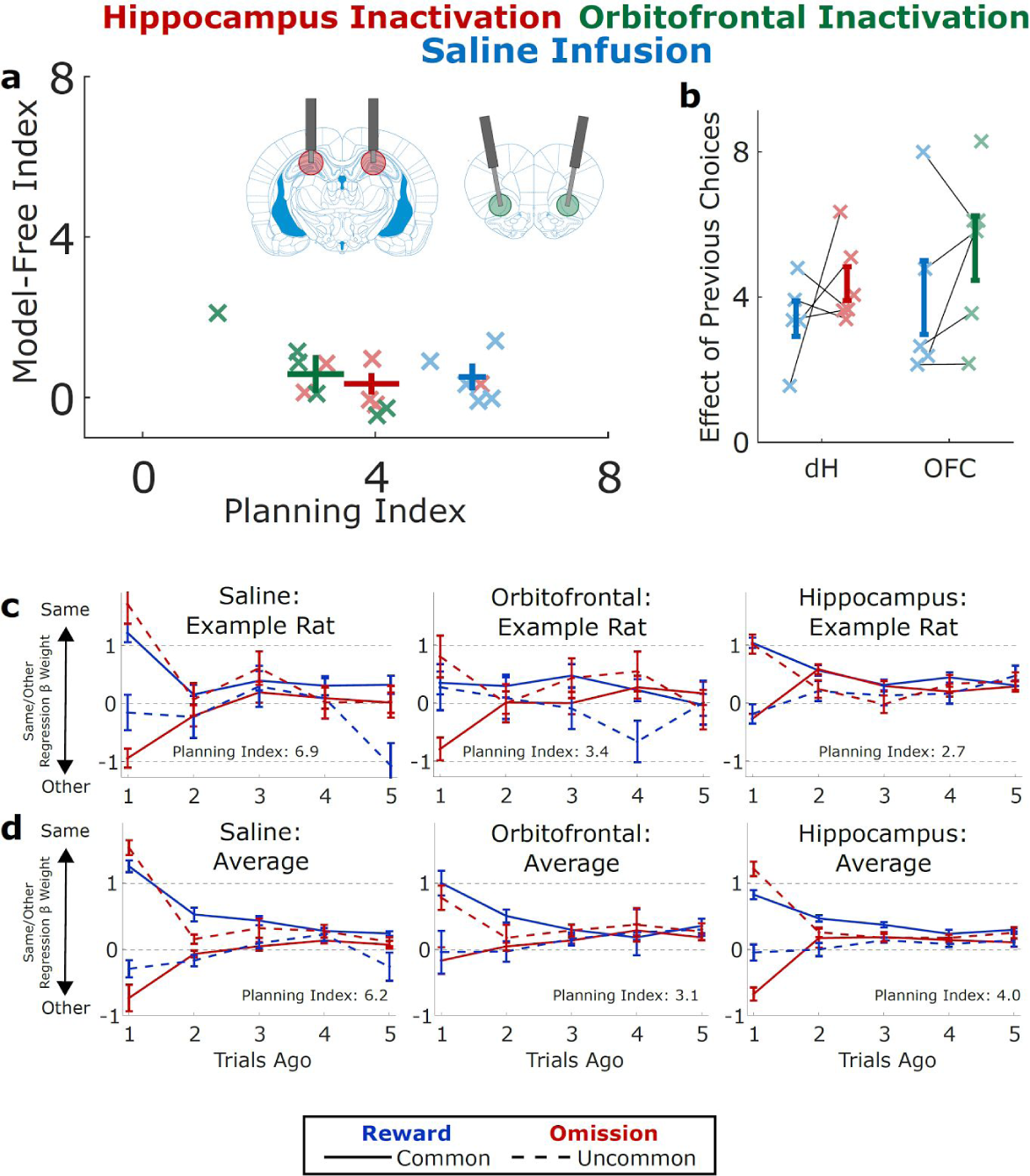
Effects of Muscimol Inactivation. A) Planning index and model-free index for implanted rats performing the task on OFC inactivation sessions (green), dH inactivation sessions (red) and pooled saline infusions (blue; pooled for display). Inactivation of either region significantly decreases the planning index. Error bars show mean across rats and standard error. B) Main effect of past choice on future choice during the same sessions (saline session unpooled). Inactivation has no significant effect on this measure. Error bars show mean across rats and standard error. C) Results of the same/other regression analysis applied to data from an example rat on saline sessions (left), OFC infusions (middle), and dH infusions (right). D) Average over all rats of the results of the same/other regression analysis.

To determine the extent to which these muscimol-induced behavioral changes were specific to planning, we applied our mixture-of-agents model to the inactivation datasets (Figure 5A; Methods, *Modeling Inactivation Data*). To make the most efficient use of our data, we adopted a hierarchical modeling approach^43–45^, simultaneously estimating parameters for both each rat individually as well as the population of rats as a whole. For each rat, we estimated the mixture-of-agents model parameters (*β*_plan_, *α*_plan_, *β*_np_, *β*_persev_, and *β*_bias_) for both control and inactivation sessions. For the population, we estimated the distribution of each of the rat-level parameters across animals, as well as the effect of inactivation on each parameter. Our population-level parameters were subject to weakly informative priors^44^ embodying the belief that infusion would most likely have little or no effect on behavioral parameters, and that any effect was equally likely to be positive or negative. To perform Bayesian inference with this model, we conditioned it on the observed datasets and used Hamiltonian Markov Chain Monte Carlo to estimate the posterior distribution jointly over all model parameters (see Methods, *Modeling Inactivation Data*; Figures S8 and S9). We summarize this distribution by reporting the median over each parameter, taking this as our estimate for that parameter. Estimates for parameters governing behavior on control sessions were similar to those produced by fitting the model to unimplanted rats (compare Figure. 5B and 4C). Estimates for parameters governing the effect of inactivation on performance suggested large and consistent effects on the planning parameter *β*_plan_, with weak and/or inconsistent effects on other behavioral components. To test whether inactivation produced significant effects on behavior that generalize to the population of rats, we compute for each population-level parameter the fraction of the posterior in which that parameter has the opposite sign as the median – the Bayesian analogue of a p-value. We found that this value was small only for the parameter corresponding to the planning weight (*β*_plan_; OFC, p = 0.01; dH, p = 0.01), and was large for all other parameters (all p > 0.1). To determine whether this was robust to tradeoff in parameter estimates between *β*_plan_ and other parameters, we inspected plots of the density of posterior samples as a function of several parameters at once. Figure 5C shows a projection of this multidimensional density onto axes that represent the change in *β*_plan_ (planning agent’s weight) and the change in *α*_plan_ (planning agent’s learning rate) due to the infusion. We found that no infusion-induced change in *α*_plan_ would allow a good fit to the data without a substantial reduction in the *β*_plan_ parameter (all of the significant density is below the “effect on *β*_plan_ = 0” axis). We find similar robustness with respect to the other population-level parameters (Figure 10).

To test the hypothesis that the effects of inactivation were specific to planning, rather than a more general effect on learning or memory, we constructed several variants of our model and compared them to one another using cross-validation. The first of these was a model in which inactivation was constrained to have equal effect on *β*_*plan*_, *β*_*np*_, and *β*_*persev*_, scaling each toward zero by an equal factor, to simulate a global effect of inactivation on memory. The second was a model in which inactivation affected only the influence of outcomes which occurred two or more trials in the past, to simulate an effect specifically on memory for more remote past events. The third was a combination of these two, allowing inactivation to have different effects on the recent and the remote past, but constraining it to affect all agents equally. We found that in all cases model comparison strongly dispreferred these alternative models, favoring a model in which inactivation has different effects on different components of behavior (log posterior predictive ratio of 42, 56, and 47 for OFC in the first, second, and third alternative models; lppr of 26, 43, and 26 for dH; see *Methods, Inactivation Model Comparison*). Taken together, these findings indicate that both OFC and dH play particular roles in supporting particular behavioral patterns, and that both play a specific role in model-based planning behavior. We find no evidence that either region plays a consistent role in supporting any behavioral component other than planning.

**Figure 5:**
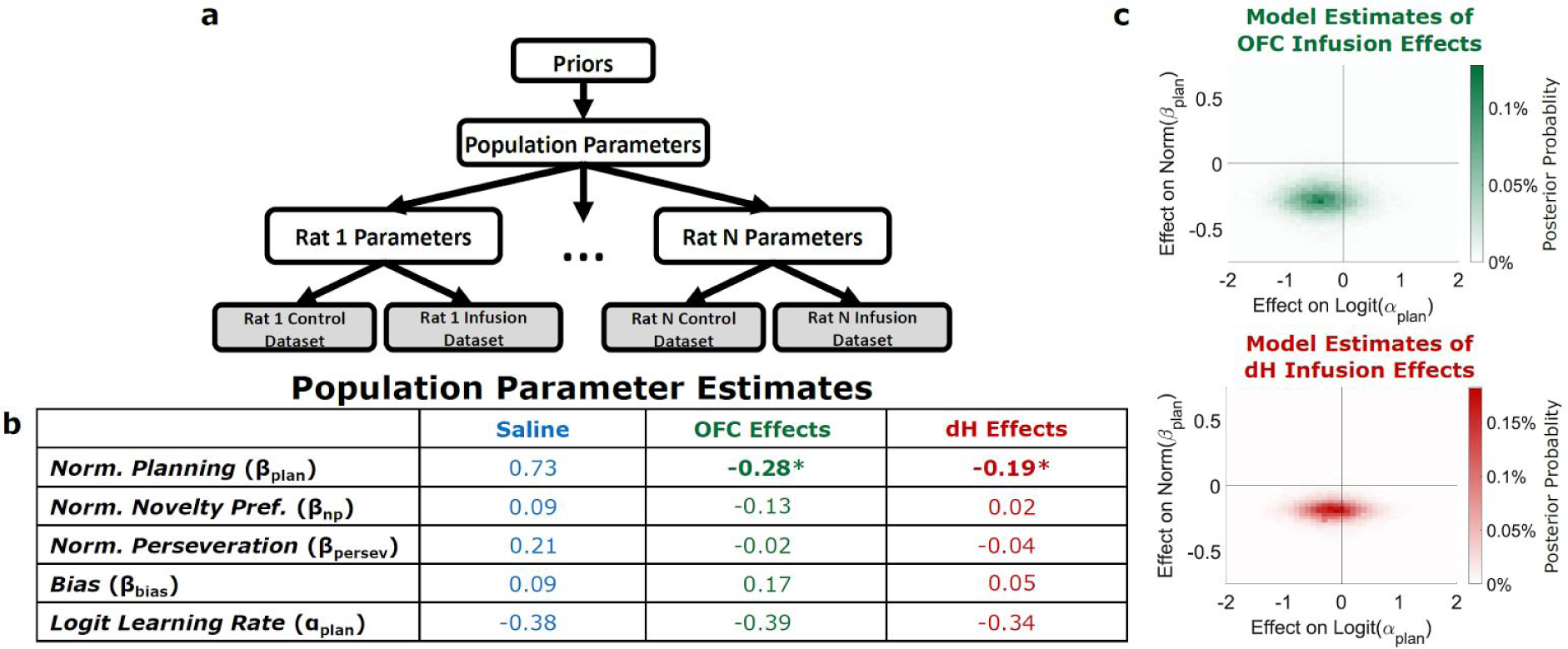
Effects of Muscimol Inactivation on Mixture Model Fits. A) Schematic showing hierarchical Bayesian framework for using the agent model for parameter estimation. Each rat is characterized by a set of control parameters governing performance in saline sessions, as well as a set of infusion effect parameters governing the change in behavior following infusion. The population of rats is characterized by the means and standard deviations of each of the rat-level parameters. These population parameters are subject to weakly informative priors. B) Parameter estimates produced by the hierarchical Bayesian model for population parameters. Column one (blue) shows parameters governing behavior on saline sessions. Columns two and three (orange and red) show parameters governing the change in performance due to OFC or dH inactivation. In columns two and three, asterisks indicate parameters for which 95% or more of the posterior distribution shares the same sign. C) Posterior belief distributions produced by the model over the parameters governing the effect of inactivation on planning weight (*β*_plan_) and learning rate(*α*_plan_).

## Discussion

We report the first successful adaptation of the two-step task – a repeated-trial, multi-step decision task widely used in human research – to rodents. This development, along with parallel efforts in other labs (Miranda, Malalasekera, Dayan, & Kennerly, 2013, *Society for Neuroscience Abstracts*; Akam, Dayan, & Costa, 2013, *Cosyne Abstracts*; Groman, Chen, Smith, Lee, & Taylor, 2014, *Society for Neuroscience Abstract*s; Hasz & Redish, 2016, *Society for Neuroscience Abstracts*), provides a broadly applicable tool for investigating the neural mechanisms of planning. While previously existing planning tasks for rodents are well-suited to identifying the neural structures involved, and have the advantage of exposing for study the process of model learning itself, the two-step task provides complementary advantages by eliciting many planned decisions in each behavioral session, opening the door to a wide variety of new experimental designs. These designs include those employing neural recordings to characterize the neural correlates of planning, as well as those, like ours, employing trial-by-trial analysis to separately quantify the relative influence of planning vs. other behavioral strategies.

Analysis of choice behavior on our task reveals a dominant role for model-based planning. Our rats also reveal knowledge of action-outcome contingencies in their movement times (Figure S3), making it unlikely that they are using any non-planning strategy, including one which might use an alternative state space to allow it to mimic model-based choice^46^ (see supplementary discussion for details on such strategies). Strikingly, our analysis reveals little or no role for model-free reinforcement learning, in contrast with the performance of human subjects on the same task^10,47–49^. One possible reason for this is the extensive experience our rat subjects have with the task – human subjects given several sessions of training tend, like our rats, to adopt a predominantly model-based strategy^50^. These data stand in tension with theoretical accounts suggesting that model-based control is a slower, more costly, or less reliable alternative to model-free, and should be avoided in situations where it does not lead to a meaningful increase in reward rates^5,51^. They are in accord with data showing that human subjects adopt model-based strategies even when this does not result in an increase in reward rate^52^. Together, these data may suggest that model-based control may be a default decision-making strategy adopted in the face of complex environments.

We found that reversible inactivation of orbitofrontal cortex selectively impaired model-based choice, consistent with previous work indicating causal roles for this region in model-based control in rodents^36–39^ and monkeys^53,54^, as well as theoretical accounts positing a role for this structure in model-based processing and economic choice^55–57^. That we observe similar effects in the rat two-step task is an important validation of this behavior as an assay of planning in the rat. It is important to note that not all accounts of OFC’s role in model-based processing are consistent with a causal role in instrumental choice^58^. Our findings here are therefore not merely confirmatory, but also help adjudicate between competing accounts of orbitofrontal function.

Inactivation of dorsal hippocampus also selectively impaired model-based control, leaving other behavioral patterns unchanged. This finding offers the first direct causal demonstration of a long-hypothesized role for hippocampus in model-based planning behavior. Long-standing theories of hippocampal function^14^ hold that the hippocampus represents a “cognitive map” of physical space, including the spatial relationships between objects and obstacles, and that this map is used in navigational planning. Classic causal data indicate that hippocampus is necessary for tasks that require navigation^15,16^, but do not speak to the question of its involvement specifically in planning. Such data are consistent with theoretical accounts in which hippocampus provides access to abstract spatial state information (i.e., location) as well as abstract spatial actions (e.g. ‘run south,’ independent of present orientation)^59^. This information might be used either by a strategy based on action-outcome associations (i.e., a planning strategy), or on stimulus-response associations (a model-free strategy). An example of this comes from experiments using the elevated plus-maze^15^, in which a rat with an intact hippocampus might adopt a strategy of running south at the intersection, independent of starting location, either because it knows that this action will lead to a particular location in the maze (planning) or because it has learned a stimulus-response mapping between this location and this spatial action. A related literature argues that the hippocampus is important for working memory^60^, citing hippocampal impairments in tasks such as delayed alternation^61^ and foraging in radial arm mazes^62,63^, in which decisions must be made on the basis of recent past events. Impairments on these tasks are consistent both with accounts in which information about the recent past is used in model-free learning (i.e. generalized stimulus-response learning in which the “stimulus” might be a memory) as well as with accounts in which it supports action-outcome planning in particular. Here, we address the action-outcome question directly, finding a role for hippocampus specifically in planning, but not other behavioral strategies. Importantly, we find that our data are *less* well explained by models in which inactivation impairs memory in general, rather than planning specifically. This indicates that if the hippocampus’ role in our task is to support memory, then this is a particular type of memory that is specifically accessible for the purposes of planning. More generally, while our data demonstrate a deficit that is specific to planning at the level of behavior, it remains unknown whether this is because the hippocampus performs computations that are integral to the planning process per se (i.e. actively using an action-outcome model to inform choice), or instead performs another function that is specifically necessary to support planning (e.g. remembering the model of actions-to-outcomes).

Our results are in accord with theoretical accounts which posit a role for the hippocampus in planning^22–25^, but stand in tension with data from classic causal experiments. These experiments have demonstrated intact action-outcome behaviors following hippocampal damage in a variety of spatial and non-spatial assays. One prominent example is latent learning, in which an animal that has previously been exposed to a physical maze learns to navigate a particular path through that maze more quickly than a naive animal – whether or not it has an intact hippocampus^26,27,31^. Hippocampal damage also has no impact on classic assays of an animal’s ability to infer causal structure in the world, including contingency degradation, outcome devaluation, and sensory preconditioning^28–30^. A comparison of these assays to our behavior reveals one potentially key difference: only the two-step task requires the chaining together of multiple action-outcome associations. Outcome devaluation, for example, requires one A-O association (e.g. lever–food), as well as the evaluation of an outcome (food–utility). Our task requires two A-O associations (e.g. top-left poke – bottom-right port lights; bottom-right poke – water) as well an evaluation (water–utility). This difference suggests one possible resolution: perhaps the hippocampus is necessary specifically in cases where planning requires linking actions to outcomes over multiple steps. This function may be related to the known causal role of hippocampus in relational memory tasks^32,33^, which require chaining together multiple stimulus-stimulus associations. It may also be related to data indicating a role in second-order classical conditioning^64^ and as well as in first-order conditioning specifically when CS and US are separated by a delay^65,66^. Future work should investigate whether it is indeed the multi-step nature of the two-step task, rather than some other feature, that renders it hippocampally dependent.

Another contentious question about the role of hippocampus regards the extent to which it is specialized for spatial navigation^67–69^ as opposed to playing some more general role in cognition^70–73^. While performing the two-step task does require moving through space, the key relationships necessary for planning on this task are non-spatial, namely the causal relationships linking first-step choice to second-step outcome (the spatial geometry of which was counterbalanced across animals). Once the first-step choice was made, lights in each subsequent port guided the animal through the remainder of the trial – apart from the single initial left/right choice, no navigation or knowledge of spatial relationships was necessary. Taken together with the literature, our results suggest that multi-step planning specifically may depend on the hippocampus, in the service of both navigation and other behaviors.

Model-based planning is a process that requires multiple computations. Importantly, our results do not reveal the particular causal role within the model-based system that is played by either hippocampus or OFC. An important question which remains open is whether these regions perform computations involved in the planning process per se (i.e. actively using an action-outcome model to inform choice), or instead perform computations which are specifically necessary to support planning (e.g. planning-specific forms of learning or memory). It is our hope that future studies employing the rat two-step task, perhaps in concert with electrophysiology and/or optogenetics, will be able to shed light on these and other important questions about the neural mechanisms of planning.

## Acknowledgements

We thank Jeff Erlich, Chuck Kopec, C. Ann Duan, Tim Hanks, and Anna Begelfer for training KJM in the techniques necessary to carry out these experiments, as well as for comments and advice on the project. We thank Nathaniel Daw, Ilana Witten, Yael Niv, Bob Wilson, Thomas Akam, Athena Akrami, and Alec Solway for comments and advice on the project, and we thank Jovanna Teran, Klaus Osorio, Adrian Sirko, Richard LaTourette, Lillianne Teachen, and Samantha Stein for assistance carrying out behavioral experiments. We especially thank Thomas Akam for suggestions on the physical layout of the behavior box and other experimental details. We thank Aaron Bornstein, Ben Scott, Alex Piet, and Lindsey Hunter for helpful comments on the manuscript. KJM was supported by training grant NIH T-32 MH065214, and by a Harold W. Dodd fellowship from Princeton University.

## Author Contributions

KJM, MMB, and CDB conceived the project. KJM designed and carried out the experiments and the data analysis, with supervision from MMB and CDB. KJM, MMB, and CDB wrote the paper, starting from an initial draft by KJM.

## Methods

### Subjects

All subjects were adult male Long-Evans rats (Taconic Biosciences, NY), placed on a restricted water schedule to motivate them to work for water rewards. Some rats were housed on a reverse 12-hour light cycle, and others on a normal light cycle – in all cases, rats were trained during the dark phase of their cycle. Rats were pair housed during behavioral training and then single housed after being implanted with cannula. Animal use procedures were approved by the Princeton University Institutional Animal Care and Use Committee and carried out in accordance with NIH standards. One infusion rat was removed from study before completion due to health reasons – this rat did not complete any saline sessions.

The number of animals used in the inactivation experiment was determined informally by comparison to similar previous studies and by resources available. Particular animals were selected for inclusion informally – they were the first three in each transition probability condition to complete training on the present version of the task, with high trial counts per session. Example animals (figures 2c 3a, and 4c) were selected on the basis of cleanly demonstrating effects that were consistent in the population. Corresponding plots for all animals can be found in supplemental figures S4 and S6.

### Behavioral Apparatus

Rats performed the task in custom behavioral chambers (Island Motion, NY) located inside sound- and light-attenuated boxes (Coulborn Instruments, PA). Each chamber was outfitted with six “nose ports” arranged in two rows of three, and with a pair of speakers for delivering auditory stimuli. Each nose port contained a white light emitting diode (LED) for delivering visual stimuli, as well as an infrared LED and infrared phototransistor for detecting rats’ entries into the port. The left and right ports in the bottom row also contained sipper tubes for delivering water rewards. Rats were placed into and removed from training chambers by technicians blind to the experiment being run.

### Training Pipeline

Here, we outline a procedure suitable for efficiently training naive rats on the two-step task. Automated code for training rats using this pipeline via the bControl behavioral control system can be downloaded from the Brody lab website.

*Phase I: Sipper Tube Familiarization.* In this phase, rats become familiar with the experimental apparatus, and learn to approach the reward ports when they illuminate. Trials begin with the illumination of the LED in one of the two reward ports, and reward is delivered upon port entry. Training in this phase continues until the rat is completing an average of 200 or more trials per day.

*Phase II: Trial Structure Familiarization.* In this phase, rats must complete all four actions of the complete task, with rewards delivered on each trial. Trials begin with the illumination of the LED in the top center port, which the rat must enter. Upon entry, one of the side ports (chosen randomly by the computer) will illuminate, and the rat must enter it. Once the rat does this, the LED in the bottom center port illuminates, and a sound begins to play indicating which of the bottom side ports will ultimately be illuminated (according to the 80%/20% transition probabilities for that rat). The rat must enter the lit bottom center port, which will cause the appropriate bottom side port to illuminate. Upon entry into this side port, the rat receives a reward on every trial. For rats in the “congruent” condition, the reward port available will be on the same side as the choice port selected 80% of the time, while for rats in the “incongruent” condition, ports will match in this way 20% of the time. “Violation trials” occur whenever the rat enters a port that is not illuminated, and result in a five second timeout and an aversive white noise sound. Training in this phase continues until the rat is completing an average of 200 or more trials per day with a rate of violation trials less than 5%.

*Phase IIIa: Performance-Triggered Flips.* In this phase, probabilistic dynamic rewards are introduced, and rats must learn to choose the choice port that is associated with the reward port which currently has higher reward probability. Trial structure is as in phase II, except that in 90% of trials both choice ports illuminate after the rat enters the top center port, and the rat must decide which choice port to enter. The rat then receives an auditory cue, and LED instructions to enter the bottom center port and one of the reward ports, as above. This phase consists of blocks, and in each block, one of the reward ports is “good” and the other is “bad”. If the good reward port is illuminated, the rat will receive a water reward for entering it 100% of the time. If the bad reward port is illuminated, the rat must enter it to move on to the next trial, but no water will be delivered. Which reward port is good and which is bad changes in blocks, and the change in blocks is enabled by the rat’s performance. Each block lasts a minimum of 50 trials, after this, the block switch is “enabled” if the rat has selected the choice port which leads most often to the “good” reward port on 80% of free choices in the last 50 trials. On each trial after the end is enabled, there is a 10% chance per trial that the block will actually switch, and the reward ports will flip their roles. Phase IIIa lasts until rats achieve an average of three to four block switches per session for several sessions in a row. Rats which show a decrease in trial count during this phase can often be re-motivated by using small rewards (~10% of the usual reward volume) in place of reward omissions at the “bad” port.

*Phases IIIb and IIIc.* The same as phase IIIa, except that the “good” and “bad” reward ports are rewarded 90% and 10%, respectively, in phase IIIb, and 80% and 20% of the time in phase IIIc. Block flips are triggered by the rat’s performance, as above. Each of these phases lasts until the rat achieves an average of two to three block changes per session for several sessions in a row.

*Phase IV: Final Task.* The same as phase IIIc, except that changes in block are no longer triggered by the performance of the rat, but occur stochastically. Each block has a minimum length of 10 trials, after which the block has a 2% chance of switching on each trial. In our experience, approximately 90% of rats will succeed in reaching the final task.

### Behavior Analysis

We quantify the effect of past trials and their outcomes on future decisions using a logistic regression analysis based on previous trials and their outcomes^74^. We define vectors for each of the four possible trial outcomes: common-reward (CR), common-omission (CO), uncommon-reward (UR), and uncommon-omission (UO), each taking on a value of +1 for trials of their type where the rat selected the left choice port, a value of -1 for trials of their type where the rat selected the right choice port, and a value of 0 for trials of other types. We define the following regression model:

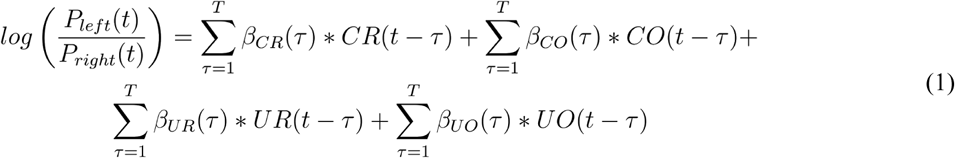

where *β_cr_*, *β_co_*, *β_ur_*, and *β_uo_* are vectors of regression weights which quantify the tendency to repeat on the next trial a choice that was made τtrials ago and resulted in the outcome of their type, and *T* is a hyperparameter governing the number of past trials used by the model to predict upcoming choice. Unless otherwise specified, *T* was set to 5 for all analyses (see Figure S2).

We expect model-free agents to show a pattern of repeating choices which lead to reward and switching away from those which lead to omissions, so we define a model-free index for a dataset as the sum of the appropriate weights from a regression model fit to that dataset:

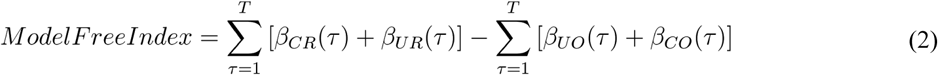

We expect that planning agents will show the opposite pattern after uncommon transition trials, since the uncommon transition from one choice is the common transition from the other choice. We define a planning index:

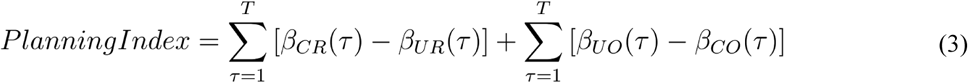

We test for significant model-free and planning indices using a one-sample t-test across rats. We test for significant differences between rats in the common-congruent and the common-incongruent conditions using a two-sample t-test.

### Behavior Models

We model our rats behavior using a mixture-of-agents approach, in which each rat’s behavior is described as resulting from the influence of a weighted average of several different “agents” implementing different behavioral strategies to solve the task. On each trial, each agent *A* computes a value, *Q*_*A*_(*a*), for each of the two available actions *a*, and the combined model makes a decision according to a weighted average of the various strategies’ values, *Q*_*total*_(*a*):

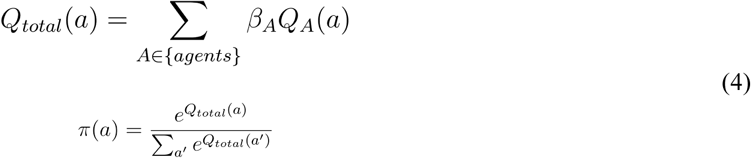

where the *β*’s are weighting parameters determining the influence of each agent, and *π(a)* is the probability that the mixture-of-agents will select action *a* on that trial. We considered models consisting of subsets of the seven following agents: model-based temporal difference learning, model-free temporal difference learning, model-based win-stay/lose switch, model-free win-stay/lose-switch, common-stay/uncommon-switch, perseveration, and bias. The “full model” consists of all of these agents, while the “reduced model” consists of four agents which were found to be sufficient to provide a good match to rat behavior. These were model-based temporal difference learning (without transition updating), novelty preference, perseveration, and bias.

*Model-Based Temporal Difference Learning.* Model-based temporal difference learning is a planning strategy, which maintains separate estimates of the probability with which each action (selecting the left or the right choice port) will lead to each outcome (the left or the right reward port becoming available), *T(a,o)*, as well as the probability, *R*_*plan*_(*o*), with which each outcome will lead to reward. This strategy assigns values to the actions by combining these probabilities to compute the expected probability with which selection of each action will ultimately lead to reward:

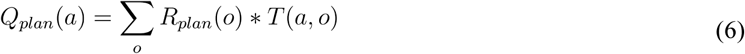

At the beginning of each session, the reward estimate *R*_*plan*_(*o*) is initialized to 0.5 for both outcomes, and the transition estimate *T(a,o)* is initialized to the true transition function for the rat being modeled (0.8 for common and 0.2 for uncommon transitions). After each trial, the reward estimate for both outcomes is updated according to

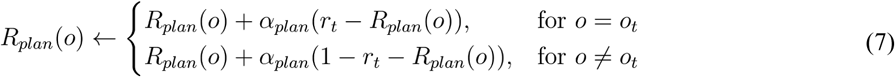

where *o*_*t*_ is the outcome that was observed on that trial, *r*_*t*_ is a binary variable indicating reward delivery, and *α*_plan_ is a learning rate parameter. The full model (but not the reduced model) also included transition learning, in which the function *T(a,o)* is updated after each outcome according to

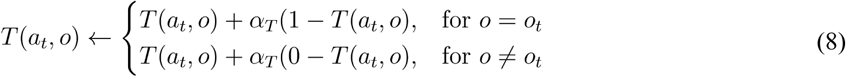

where *a*_*t*_ is the action taken, and *α*_*T*_ is a learning rate parameter.

*Model-Free Temporal Difference Learning.* Model-free temporal difference learning is a non-planning reward-based strategy. It maintains an estimate of the value of the choice ports, *Q*_*MF*_(*a*), as well as an estimate of the values of the reward ports, *R*_*MF*_(*o*). After each trial, these quantities are updated according to

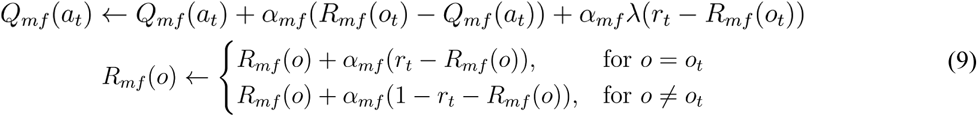

where *α*_mf_ and λ are learning rate and eligibility trace parameters affecting the update process.

*Model-Free Win-Stay/Lose Switch.* Win-stay lose-switch is a pattern that tends to repeat choices that led to rewards on the previous trial and switch away from choices that led to omissions. It calculates its values on each trial according to the following

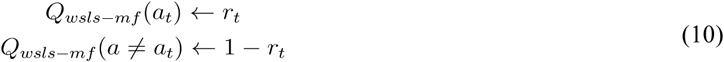

*Model-Based Win-Stay/Lose-Switch* Model-based win-stay lose switch follows the win-stay lose-switch pattern after common transition trials, but inverts it after uncommon transition trials.

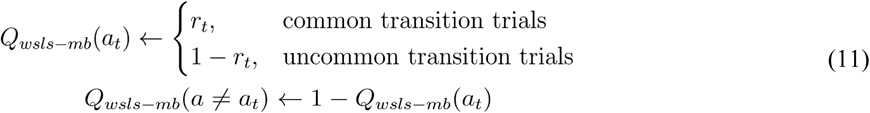

*Novelty Preference.* The novelty preference agent follows an “uncommon-stay/common switch” pattern, which tends to repeat choices when they lead to uncommon transitions on the previous trial, and to switch away from them when they lead to common transitions. Note that some rats have positive values of the *β*_np_ parameter weighting this agent (novelty preferring) while others have negative values (novelty averse; see Fig 3C):

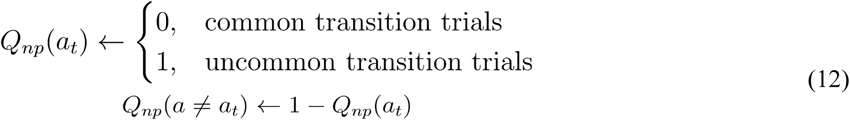

*Perseveration.* Perseveration is a pattern which tends to repeat the choice that was made on the previous trial, regardless of whether it led to a common or an uncommon transition, and regardless of whether or not it led to reward.

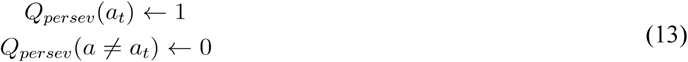

*Bias.* Bias is a pattern which tends to select the same choice port on every trial. Its value function is therefore static, with the extent and direction of the bias being governed by the magnitude and sign of this strategy’s weighting parameter *β*_*bias*_.

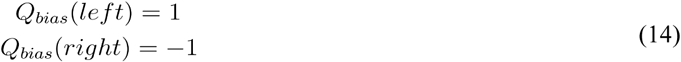

### Model Comparison and Parameter Estimation: Unimplanted Rats

We implemented the model described above using the probabilistic programming language Stan^75,76^, and performed maximum-a-posteriori fits using weakly informative priors on all parameters^44^ The prior over the weighting parameters *β* was normal with mean 0 and sd 0.5, and the prior over *α*_mf_, *α*_mb_, and λ was a beta distribution with *a*=*b*=3.

To perform model comparison, we used two-fold cross validation, dividing our dataset for each rat into even- and odd-numbered sessions, and computing the log-likelihood of each partial dataset using parameters fit to the other. For each model for each rat, we computed the “normalized cross-validated likelihood” by summing the log-likelihoods for the even- and odd-numbered sessions, dividing by the total number of trials, and exponentiating. This value can be interpreted as the average per-trial likelihood with which the model would have selected the action that the rat actually selected. We define the “reduced model” to be the full model defined above, with the parameters *β*_*MF*_, *β*_*WSLS-MF*_, *β*_*WSLS-MB*_, and *α*_*T*_ all set to zero, leaving as free parameters *β*_*plan*_, *α_plan_, β_np_, β*_*persev*_, and *β*_*bias*_ (note that *α*_*mf*_ and *λ* become undefined when *β*_*MF*_ = 0). We compared this reduced model to nine alternative models: four in which we allowed one of the fixed parameters to vary freely, four in which we fixed one of the free parameters *β*_*plan*_, *β*_*np*_, *β*_*persev*_, or *β*_*bias*_ to zero, and the full model in which all parameters are allowed to vary.

We performed parameter estimation by fitting the reduced model to the entire dataset generated by each rat (as opposed to the even/odd split used for model comparison), using maximum-a-posteriori fits under the same priors. For ease of comparison, we normalize the weighting parameters *β*_*plan*_, *β*_*np*_, and *β*_*persev*_, dividing each by the standard deviation of its agent’s associated values (*Q*_*plan*_, *Q*_*np*_, and *Q*_*persev*_) taken across trials. Since each weighting parameter affects behavior only by scaling the value output by its agent, this technique brings the weights into a common scale and facilitates interpretation of their relative magnitudes, analogous to the use of standardized coefficients in regression models.

### Synthetic Behavioral Datasets: Unimplanted Rats

To generate synthetic behavioral datasets, we took the maximum-a-posteriori estimates parameter estimates for each rat, and used the reduced model in generative mode. The model matched to each rat received the same number of trials as that rat, as well as the same sequence of reward probabilities. We used these synthetic datasets for qualitative model-checking: if the reduced model does a good job capturing patterns in behavior, applying the regression analysis to both real and synthetic datasets should yield similar results.

### Surgery

We implanted six rats with infusion cannula targeting dorsal hippocampus, orbitofrontal cortex, and prelimbic cortex, using standard stereotaxic techniques (data from PL are not reported in this paper). Anesthesia was induced using isoflurane, along with injections of ketamine and buprenorphine, the head was shaved, and the rat was placed in a stereotax (Kopf instruments) using non-puncture earbars. Lidocaine was injected subcutaneously under the scalp for local anesthesia and to reduce bleeding. An incision was made in the scalp, the skull was cleaned of tissue and bleeding was stopped. Injection cannula were mounted into guide cannula held in stereotax arms (dH & OFC: 22ga guide, 28ga injector; PL: 26ga guide, 28ga injector; Plastics One, VA), while a separate arm held a fresh sharp needle. The locations of bregma and interaural zero were measured with the tip of each injector and with the needle tip. Craniotomies were performed at each target site, and a small durotomy was made by piercing the dura with the needle. The skull was covered with a thin layer of C&B Metabond (Parkell Inc., NY), and the cannula were lowered into position one at a time. Target locations relative to bregma were AP -3.8, ML +-2.5, DV -3.1 for dorsal hippocampus, AP +3.2, ML +- 0.7, DV -3.2 for prelimbic, and AP + 3.5, ML +-2.5, DV - 5 for orbitofrontal cortex. Orbitofrontal cannula were implanted at a 10 degree lateral angle to make room for the prelimbic implant. Cannula were fixed to the skull using Absolute Dentin (Parkell Inc, NY), and each craniotomy was sealed with Kwik-Sil elastomer (World Precision Instruments, FL). One all cannula were in place, Duralay dental acrylic (Reliance Dental, IL) was applied to secure the implant. The injector was removed from each guide cannula, and replaced with a dummy cannula. Rats were treated with ketofen 24 and 48 hours post-operative, and allowed to recover for at least seven days before returning to water restriction and behavioral training.

### Inactivation Experiments

Each day of infusions, an injection system was prepared with the injection cannula for one brain region. The injection cannula was attached to a silicone tube, and both were filled with light mineral oil. A small amount of distilled water was injected into the other end of the tube to create a visible water-oil interface, and this end was attached to a Hamilton syringe (Hamilton Company, NV) filled with distilled water. This system was used to draw up and let out small volumes of muscimol solution, and inspected to ensure that it was free of air bubbles.

Rats were placed under light isoflurane anesthesia, and the dummy cannula were removed from the appropriate guide cannula. The injector was placed into the guide, and used to deliver 0.3 uL of 0.25 mg/mL muscimol solution over the course of 90 seconds. The injector was left in place for four minutes for solution to diffuse, and then the procedure was repeated in the other hemisphere. For saline control sessions, the same procedure was used, but sterile saline was infused in place of muscimol solution. The experimenter was not blind to the region (OFC, dH, PL) or substance (muscimol, saline) being infused. After the completion of the bilateral infusion, rats were taken off of isoflurane and placed back in their home cages, and allowed to recover for 30-60 minutes before being placed in the behavioral chamber to perform the task.

### Analysis of Inactivation Data

For each rat, we considered five types of sessions: OFC muscimol, dH muscimol, OFC control, dH control, and saline. Control sessions were performed the day before and the day after each infusion session, and saline sessions were pooled across OFC saline infusions and dH saline infusions (OFC musc., 18 sessions, OFC cntrl, 36 sessions, OFC sal. 6 sessions, dH musc. 33 sessions, dH cntrl, 64 sessions, dH sal. 10 sessions). Our dataset for each session consisted of up to the first 400 trials of each session in which at least 50 trials were performed. We perform the regression analysis (equation one), and compute the model-free index and planning index (equations two and three) for each dataset. To compute p-values, we performed a paired t-test across rats on the difference between muscimol and control datasets for each region, and on the difference between muscimol infusion in each region and the pooled saline infusion datasets.

### Modeling Inactivation Data

We constructed a hierarchical Bayesian version of our reduced model, using the probabilistic programming language Stan. This model considered simultaneously two datasets from each rat: an inactivation and a control dataset. Each of these datasets is modeled as the output of the reduced model (see *Behavior Models*, above), which take the five parameters *β*_*plan*_, *α*_plan_, *β*_*np*_, *β*_*persev*_, and *β*_*bias*_, giving each rat ten parameters: five for the control dataset, and five for the infusion dataset. For the purpose of the hierarchical model, we reparameterize these, characterizing each rat *R* by ten parameters organized into two vectors, 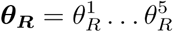 and 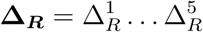 according to the following mapping:

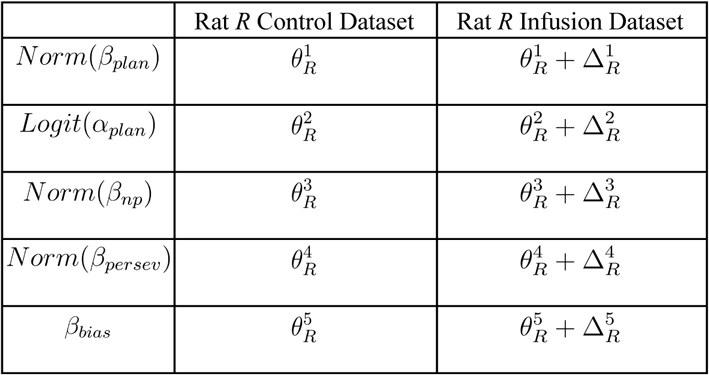

where *Norm* indicates normalization of the weight (see *Parameter Estimation*, above), and *Logit* indicates the inverse-sigmoid logit function, which transforms a parameter bounded at 0 and 1 into a parameter with support over all real numbers.

The values in **θ**_**R**_ and **Δ**_**R**_ adopted by a particular rat are modeled as draws from a gaussian distribution governed by population-level parameter vectors **θ**_**μ**_, **θ**_**σ**_, **Δ**_**μ**_, and **Δ**_**σ**_ giving the mean and standard deviation of the distribution of each of the rat-level parameters in the population:

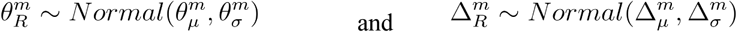

for each rat *R*, for each value of *m* indexing the various parameter vectors.

These population-level parameters are themselves modeled as draws from weakly informative prior distributions^44^ chosen to enforce reasonable scaling and ensure that all posteriors were proper:

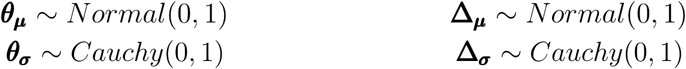

Having established this generative model, we perform inference by conditioning it on the observed datasets (control and inactivation) for each rat, and approximating the joint posterior over all parameters by drawing samples using Hamiltonian Markov Chain Monte Carlo (H-MCMC)^44,77^. To obtain estimated values for each parameter, we take the median of these samples with respect to that parameter. To test whether inactivation produced effects on behavior that were consistent at the population level, we computed a “p-value” for each parameter in **Δ**_**μ**_ given by the fraction of samples having the opposite sign as the median sample.

### Inactivation Model Comparison

We performed a series of model comparisons between models like the above and alternative models in which inactivation affected memory in general, memory for distant past trials specifically, or a combination of these. In the first alternative model, inactivation was constrained to affect equally all of the agents which depend on the history of previous trials (planning, perseveration, and novelty preference). This alternative model contains a new parameter, the “memory multiplier”, *m*, which scales the weights of these agents, in this revised version of eq. 4:

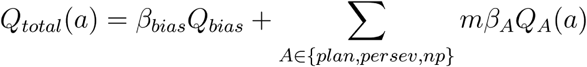

This memory multiplier is fixed to 1 for control sessions, but allowed to vary freely for each rat in infusion sessions. It has it’s own population-level mean and variance parameters, which are given weakly informative priors (see Methods, Modeling Inactivation Data). In the alternative version of the model, the *β*_*A*_ parameters are fixed between control and inactivation sessions. Since bias does not require memory, *β*_*bias*_ is allowed to vary. We implement this by fixing the parameters 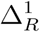 through 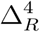 to zero for each rat R (see Methods, Modeling Inactivation Data), and allowing the effects of inactivation to be described by 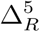 and the new parameter *m*_*R*_.

We compare this model to the model above using two-fold cross validation of H-MCMC fits. To compare these models quantitatively, we compute the log posterior predictive ratio (lppr):

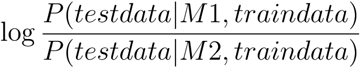

In the next model comparisons, we separate the influence of the most recent trial’s outcome from the influence of all trials further back in time. We implement this by replacing the model-based reinforcement learning agent (equations 6 & 7) with both a model-based win-stay/lose-switch agent (equation 11), and a new “lagged model-based” agent constructed by taking the value of *Q*_*plan*_ from one trial in the past and using it to guide decision-making on the current trial, so that the value of *Q*_*lagged-mb*_ used on each trial contains information about the outcomes of all past trials except the most recent one. Fits of this model therefore contain two parameters to quantify planning: *β*_*ws/ls–mb*_ for the influence of the most recent outcome, and *β*_*lagged–mb*_ for the influence of all trials further into the past.

For the second model comparison, we limit the influence of inactivation to only affect *β*_*lagged–mb*_ and *β*_*bias*_– that is, to only affect the influence of distant past trials on choice behavior, as well as choice bias. For the this model comparison, we also allow inactivation to affect the memory multiplier *m*, allowing it to have separate effects on memory for distant past trials and on memory for the immediately previous trial. We compare both of these models to a model in which inactivation can have separate effects on the each of the components of behavior. We compute the log posterior predictive ratio using leave-one-out cross validation over sessions (i.e., we compute posteriors based on all of the dataset except for one session, and compute the lppr for that session using those posteriors, then repeat for all sessions).

### Synthetic Behavioral Datasets: Inactivation Data

To generate synthetic behavioral datasets, we took the parameter estimates produced by the hierarchical model for each rat for orbitofrontal, hippocampus, and saline infusions. Parameters used for synthetic saline datasets were the average of the saline parameters produced by fitting the model to saline/hippocampus data and to saline/orbitofrontal (note that rat #6 did not complete any saline sessions – parameter estimates for this rat are still possible in the hierarchical model since they can be “filled in” based on infusion data and data from other rats). We used the reduced model in generative mode with these parameters, applying each parameter set to a simulated behavioral session consisting of 10,000 trials. We then applied the trial-history regression analysis to these synthetic datasets, and used the results for qualitative model checking, comparing them to the results of the same analysis run on the actual data.

### Code and Data Availability

All software used for behavioral training will be available on the Brody lab website. Software used for data analysis, as well as raw and processed data, are available from the authors upon request.

## Supplementary Figures

**Figure S1:**
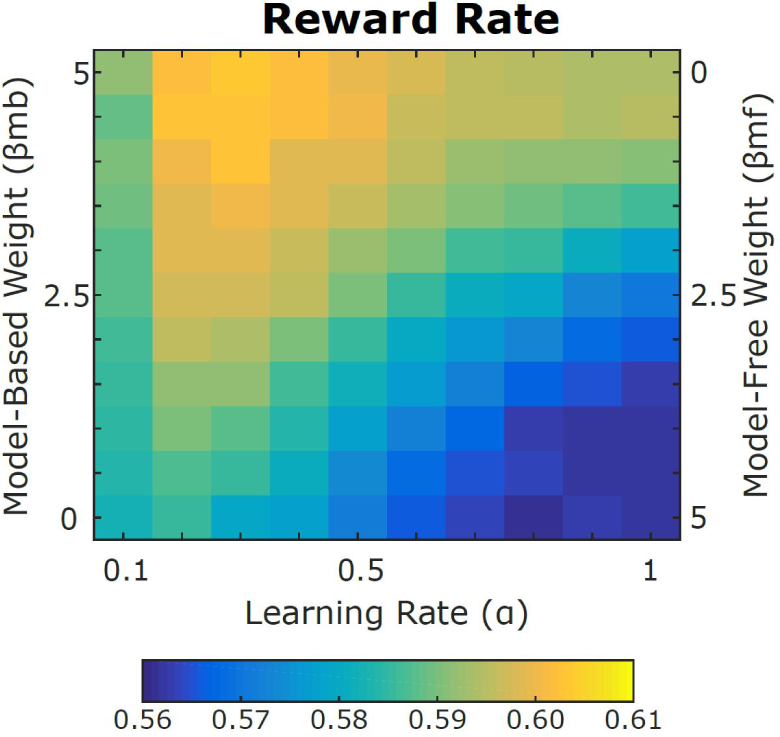
Reward Rates of Model-Based & Model-Free Agents. Reward rates achieved by synthetic datasets generated by a hybrid model-based/model-free agent. Data were generated under the constraints α_plan_=α_MF_, β_plan_+β_MF_=5, with λ, α_T_, and all other betas set to zero. The highest reward rates are achieved by purely model-based agents, but the best purely model-free agents still outperformed the average rat, earning around 58% rewards (rat’s reward rate mean: 56.8%, sem: 0.4%, std: 2%).

**Figure S2:**
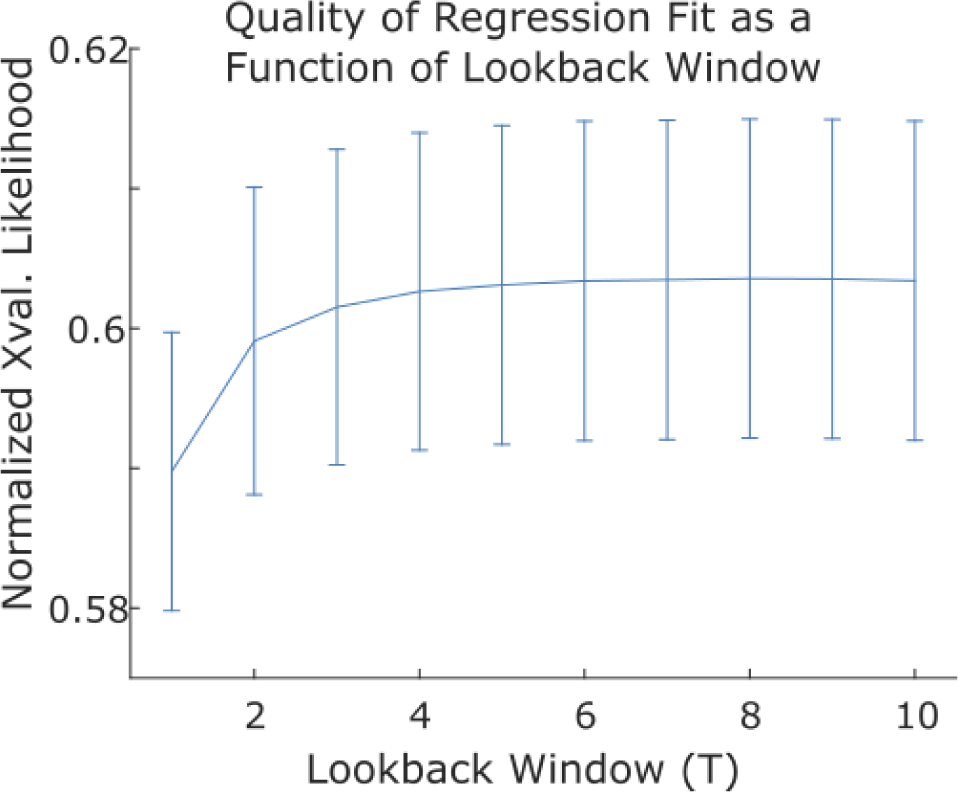
Normalized cross-validated likelihood for logistic regression models (see Methods, *Behavior Analysis*), as a function of the number of previous trials used to predict the upcoming choice. Including more than five previous trials in the model results in negligible improvements in quality of model fit.

**Figure S3:**
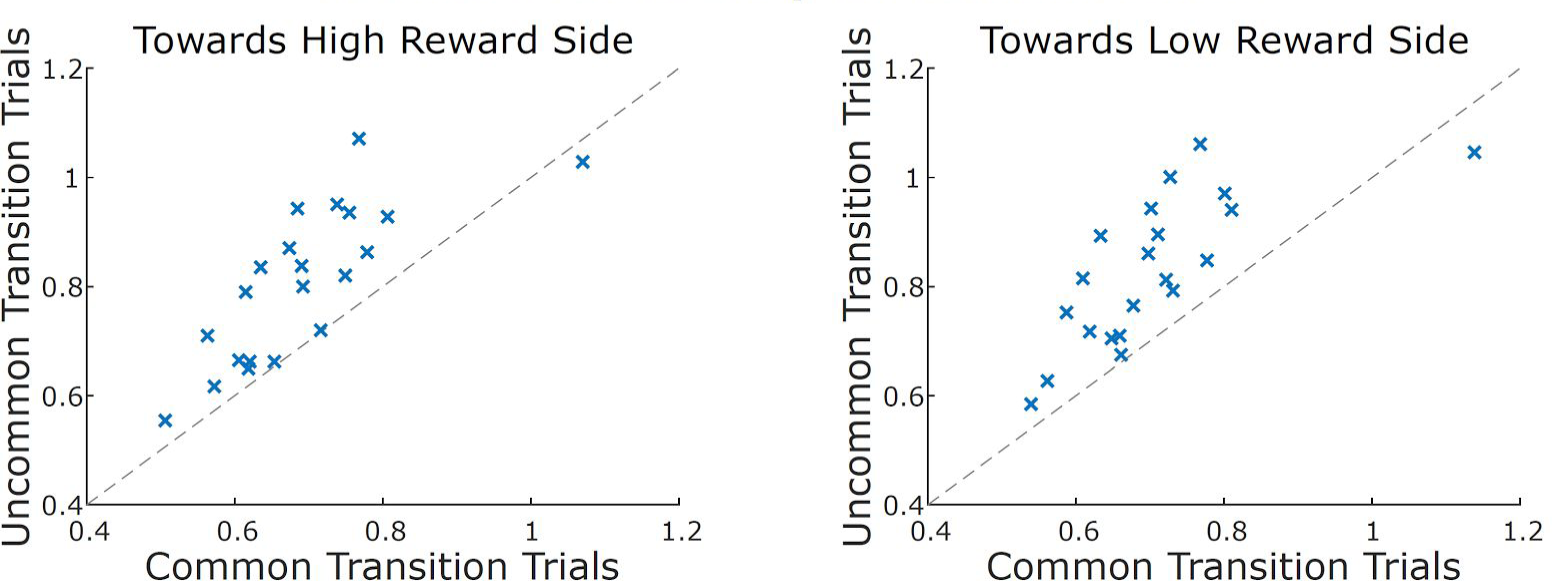
Movement times are faster following common transition trials. Median movement time, in seconds, from the bottom center port to the reward port for common and uncommon transition trials, broken down by whether the movement was towards (right panel) or away from (left panel) the port with the higher reward probability.

**Figure S4:**
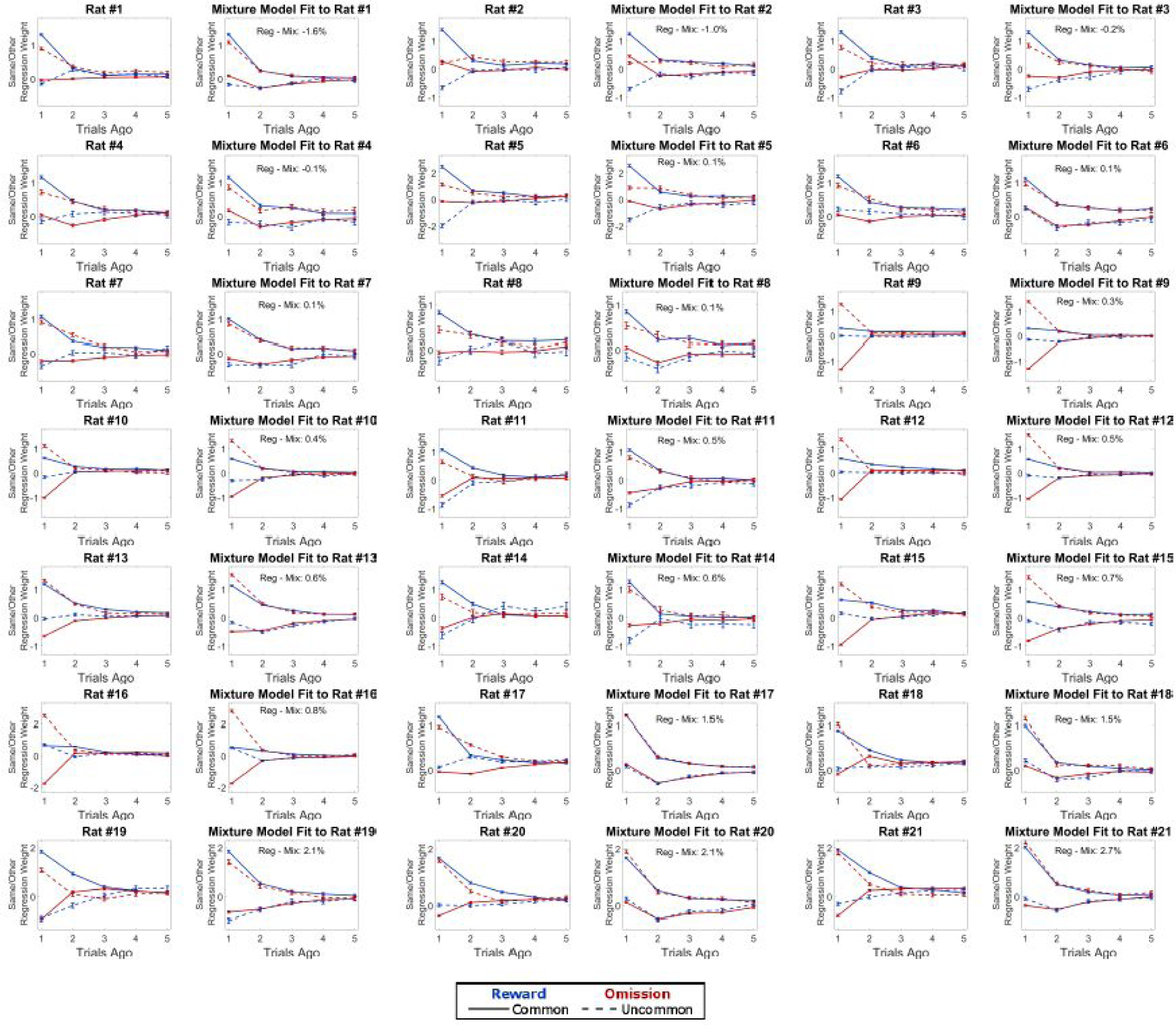
Results of logistic regression analysis applied to each rat, as well as simulated data generated from a fit of the mixture model to that rat’s dataset. Rats are ordered by the relative quality of fit of the mixture model with respect to the regression model - earlier rats datasets are better explained by the mixture model than the regression, while later rats are better explained by the regression model.

**Figure S5:**
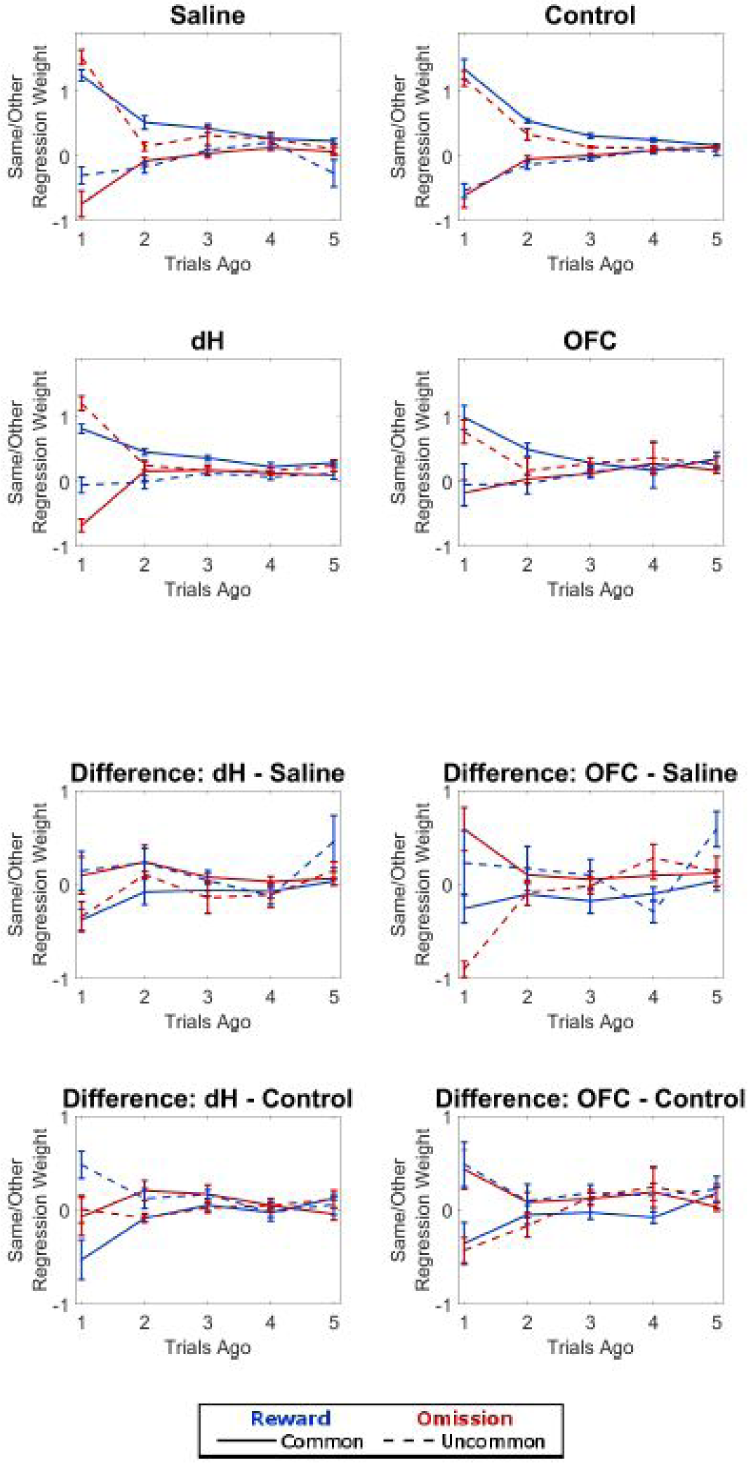
Results of logistic regression analysis applied to the inactivation dataset. Above: Regression coefficients for the Saline, Control, dH, and OFC conditions. Points are averages across rats, and error bars are standard errors. Below: Differences between regression coefficients for different conditions

**Figure S6:**
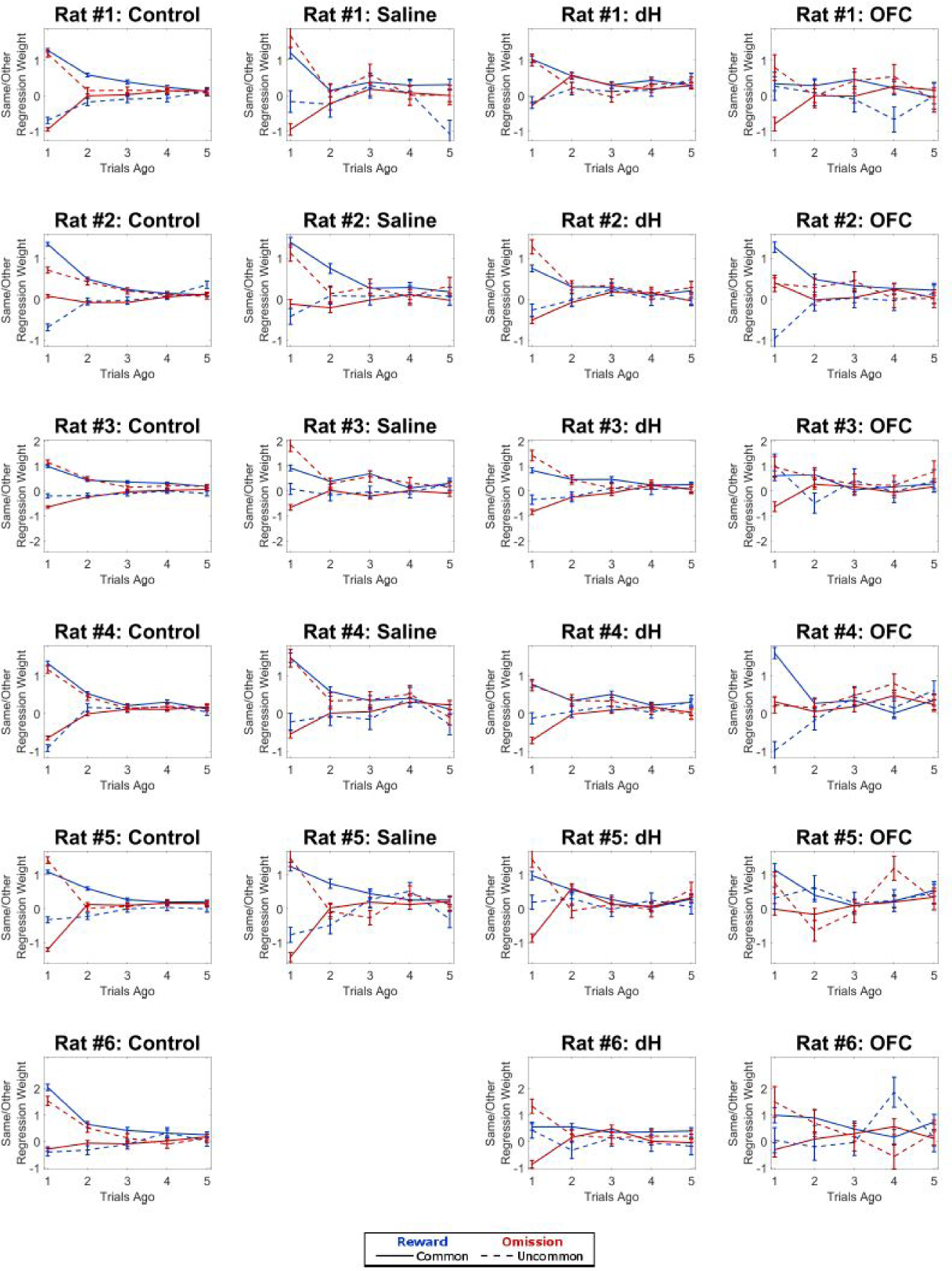
Results of logistic regression analysis applied to each rat in the inactivation dataset. Note that rat #6 did not complete any saline sessions

**Figure S7:**
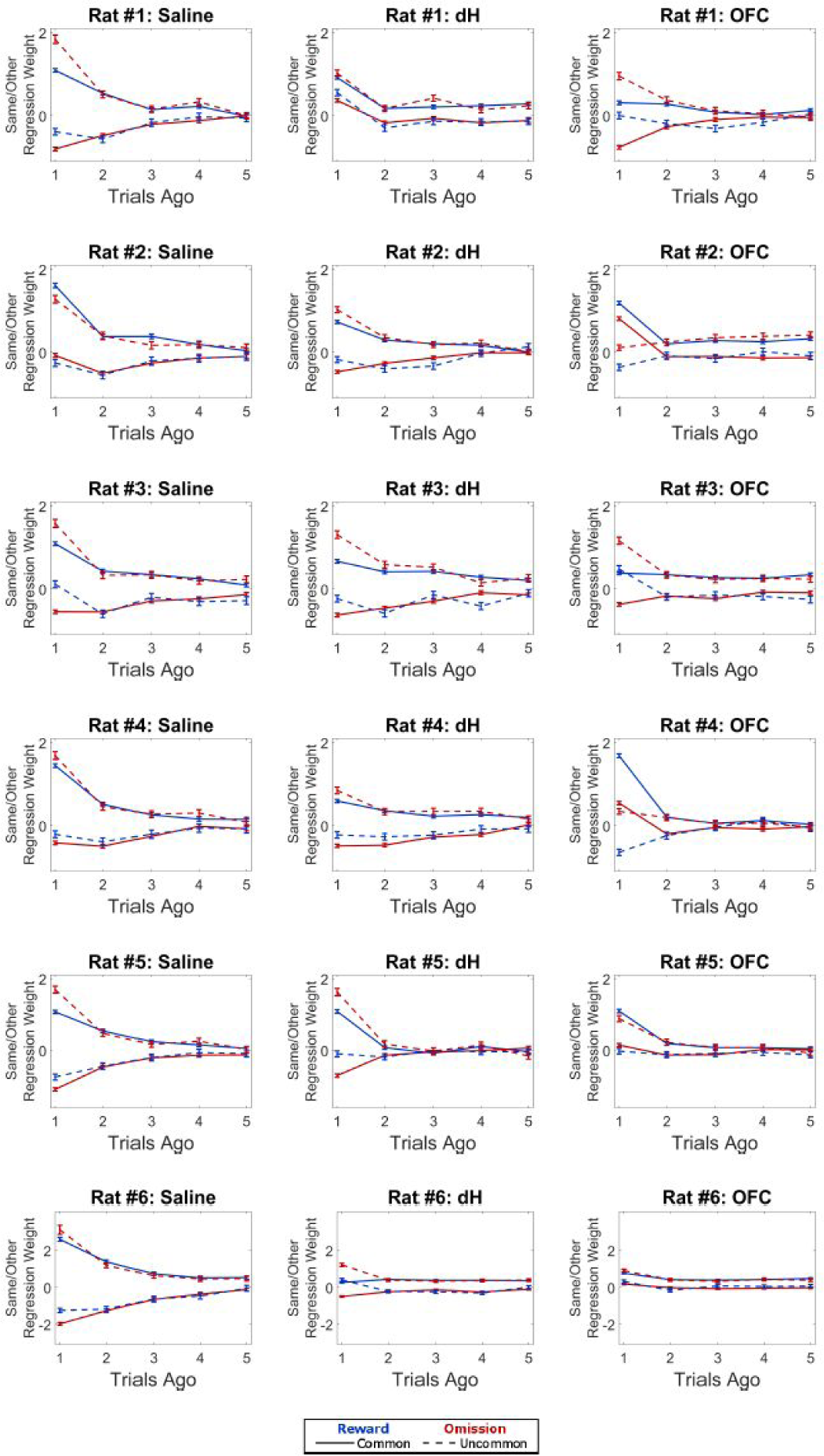
Results of logistic regression analysis applied to simulated data generated by the reduced model fit to each rat in the inactivation dataset

**Figure S8:**
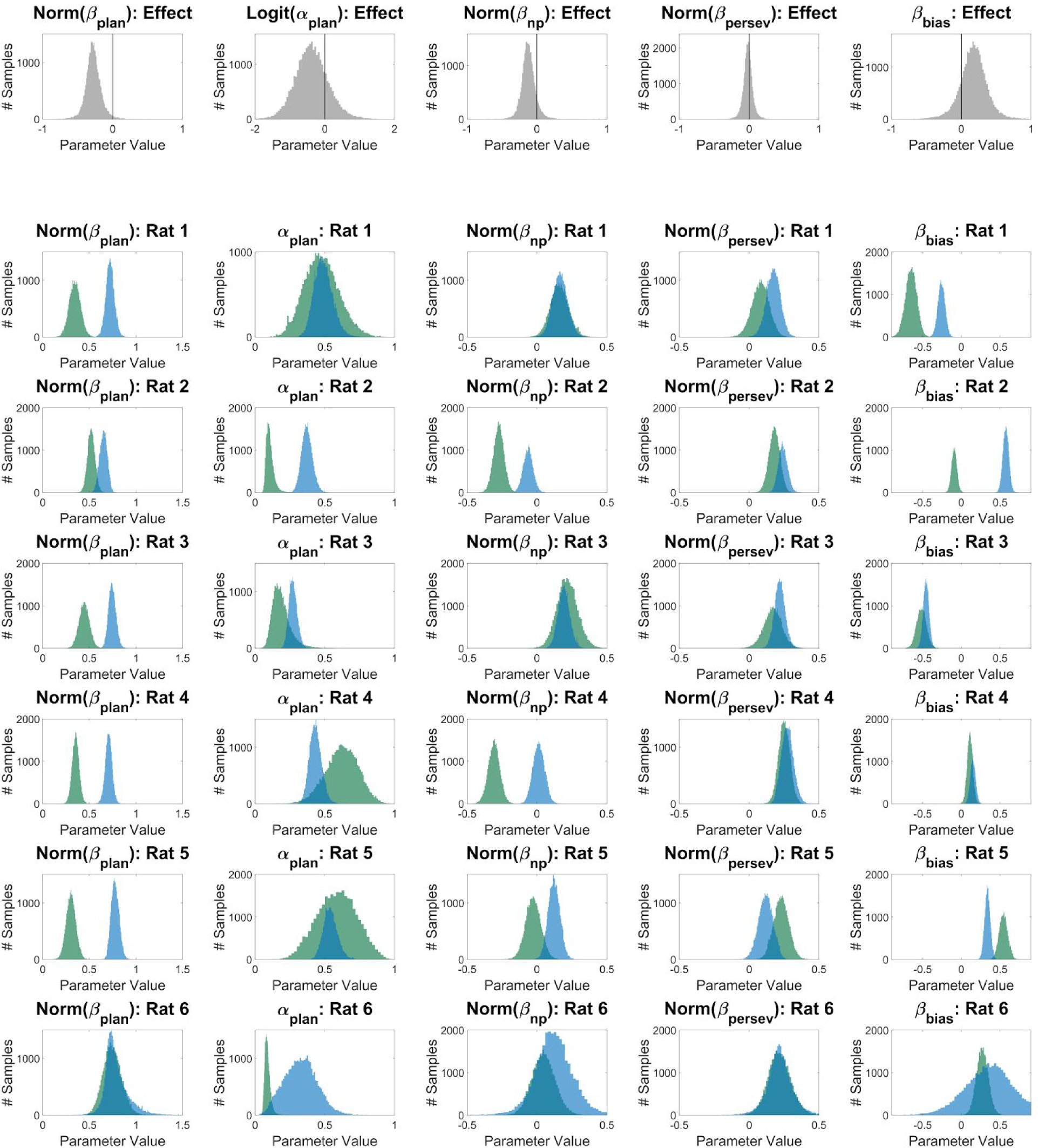
Results of fitting the multi-agent model jointly to the OFC inactivation and Saline datasets. Top Row: Posterior belief distributions over parameters governing the effect of inactivation on performance across the population. Only β_plan_ is significantly affected by the inactivation. Below: Posterior belief distributions over parameters governing behavior on OFC (green) and Saline (blue) sessions. Only β_plan_ is affected by inactivation in a way that is consistent across animals.

**Figure S9:**
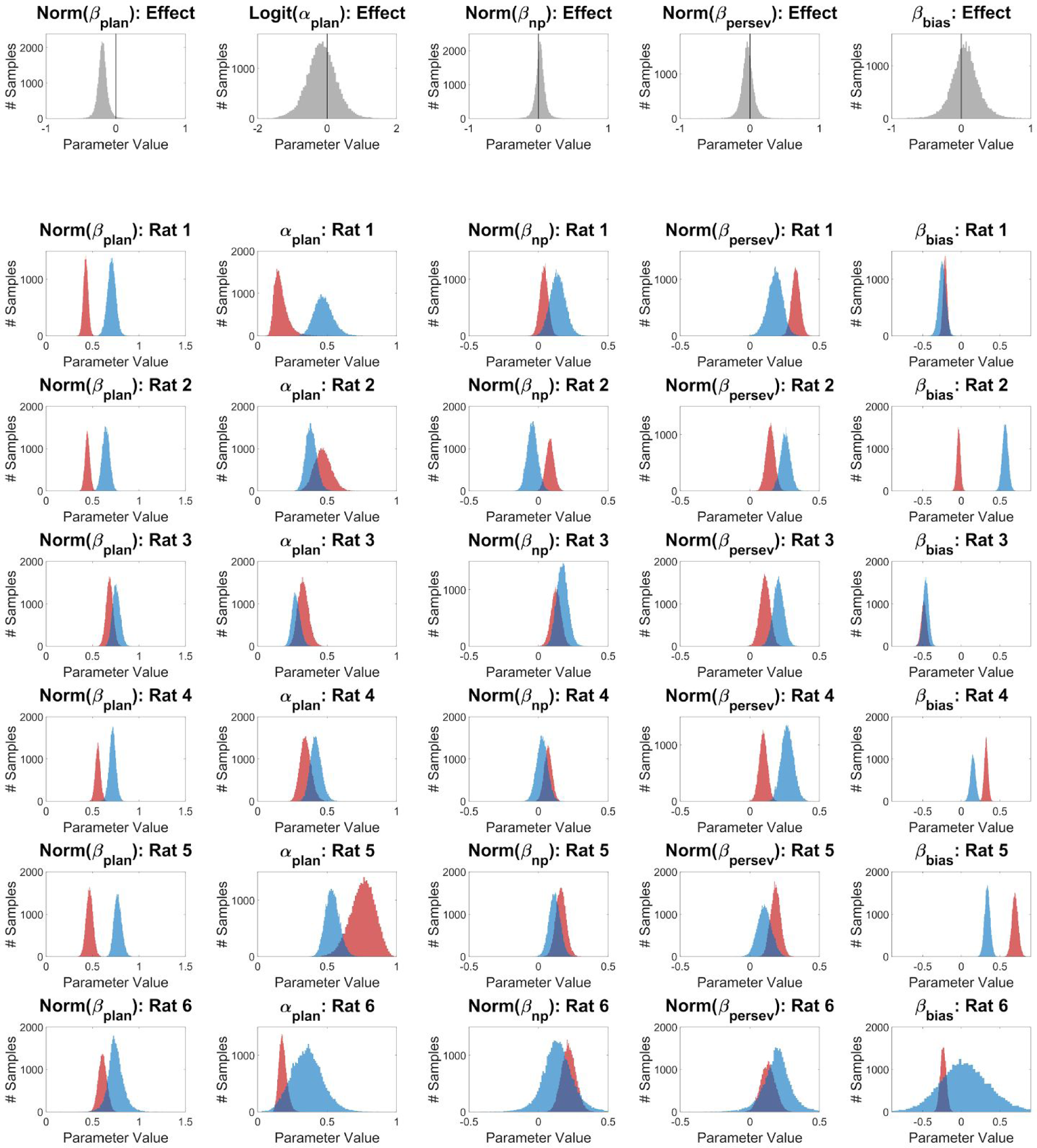
Results of fitting the multi-agent model jointly to the dH inactivation and Saline datasets. Top Row: Posterior belief distributions over parameters governing the effect of inactivation on performance across the population. Only β_plan_ is significantly affected by the inactivation. Below: Posterior belief distributions over parameters governing behavior on dH (red) and Saline (blue) sessions. Only β_plan_ is affected by inactivation in a way that is consistent across animals.

**Figure S10:**
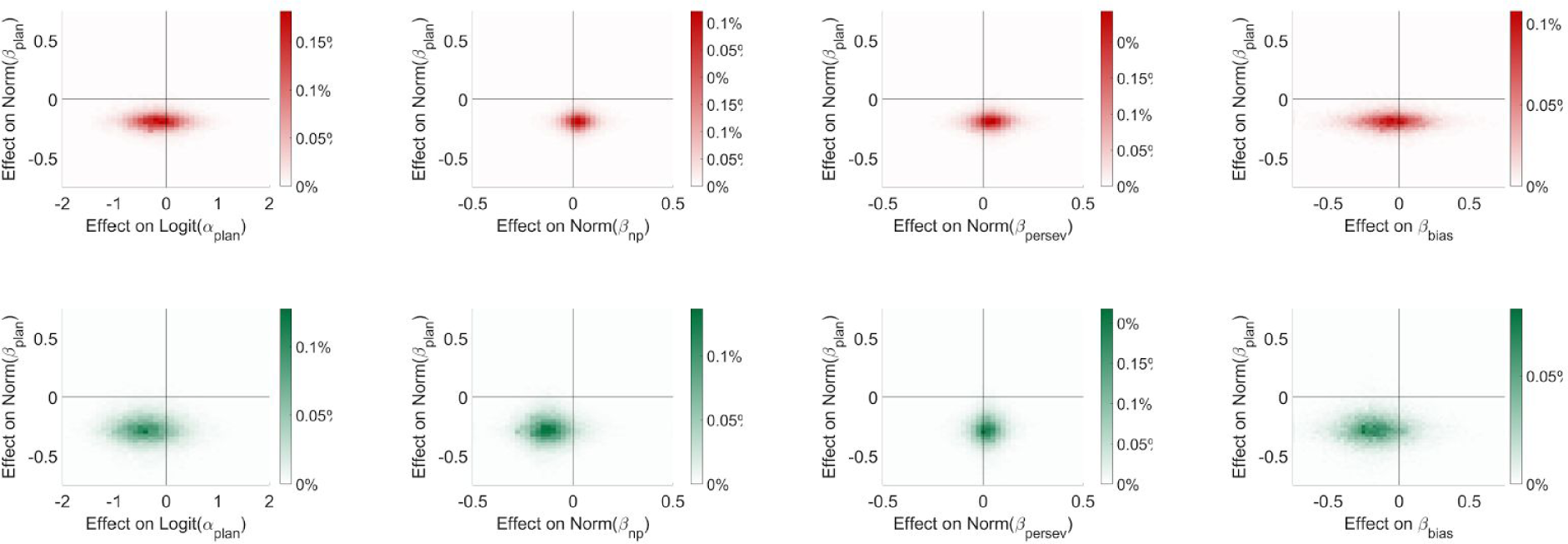
Plots of posterior density projected onto planes defined by the parameter governing change in model-based weight and other population parameters for hippocampus (above, red) and OFC (below, green) inactivation datasets.

**Figure S11:**
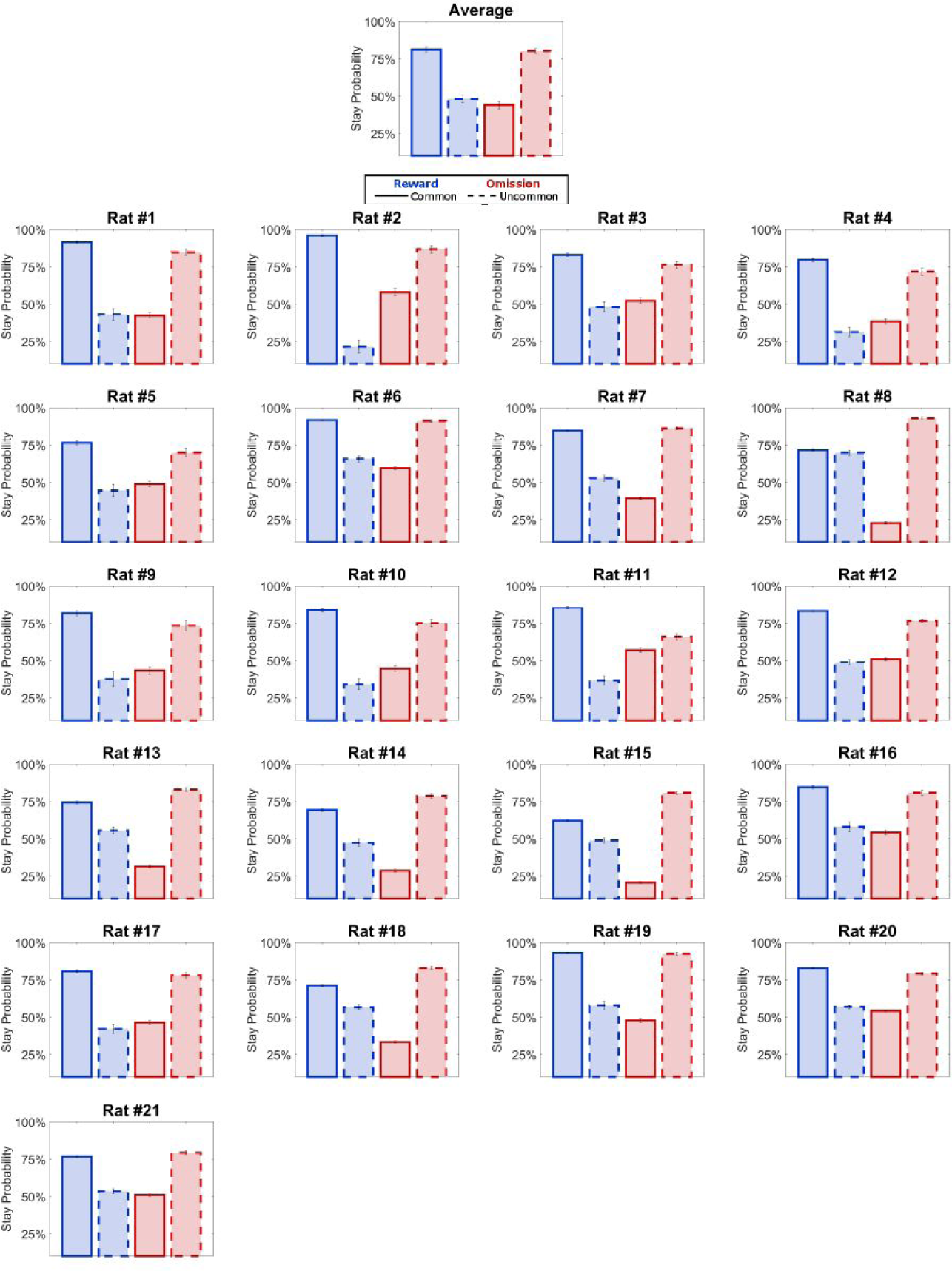
Results of one-trial-back analysis applied the behavioral dataset. Above: Average and standard error of the stay probability across rats. Below: Stay probability for each rat, with binomial 95% confidence intervals

**Figure S12:**
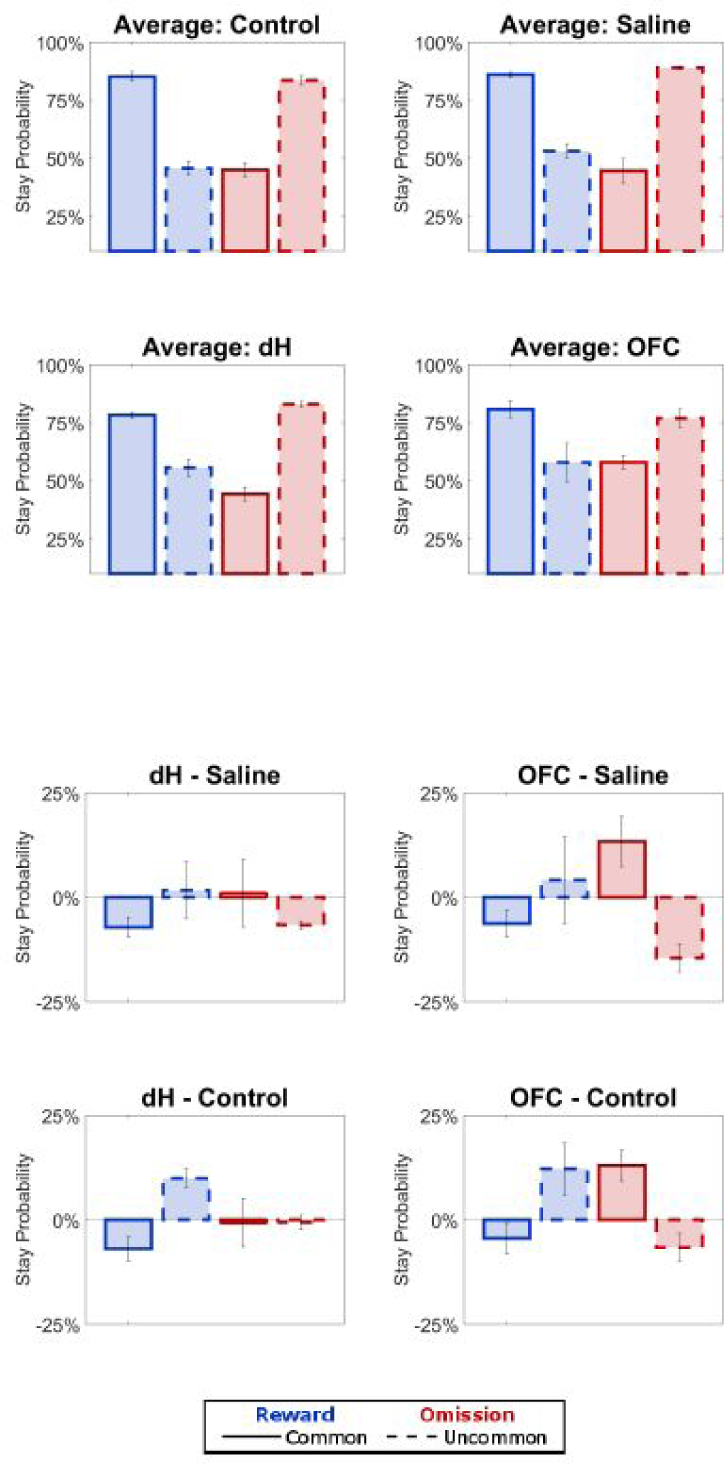
Results of one-trial-back analysis applied to the inactivation dataset. Above: Average stay probability by trial-type for the Control, dH, and OFC conditions. Bar height is the average across rats, and error bars are standard errors. Below: Differences between stay probabilities coefficients for the different conditions

**Figure S13:**
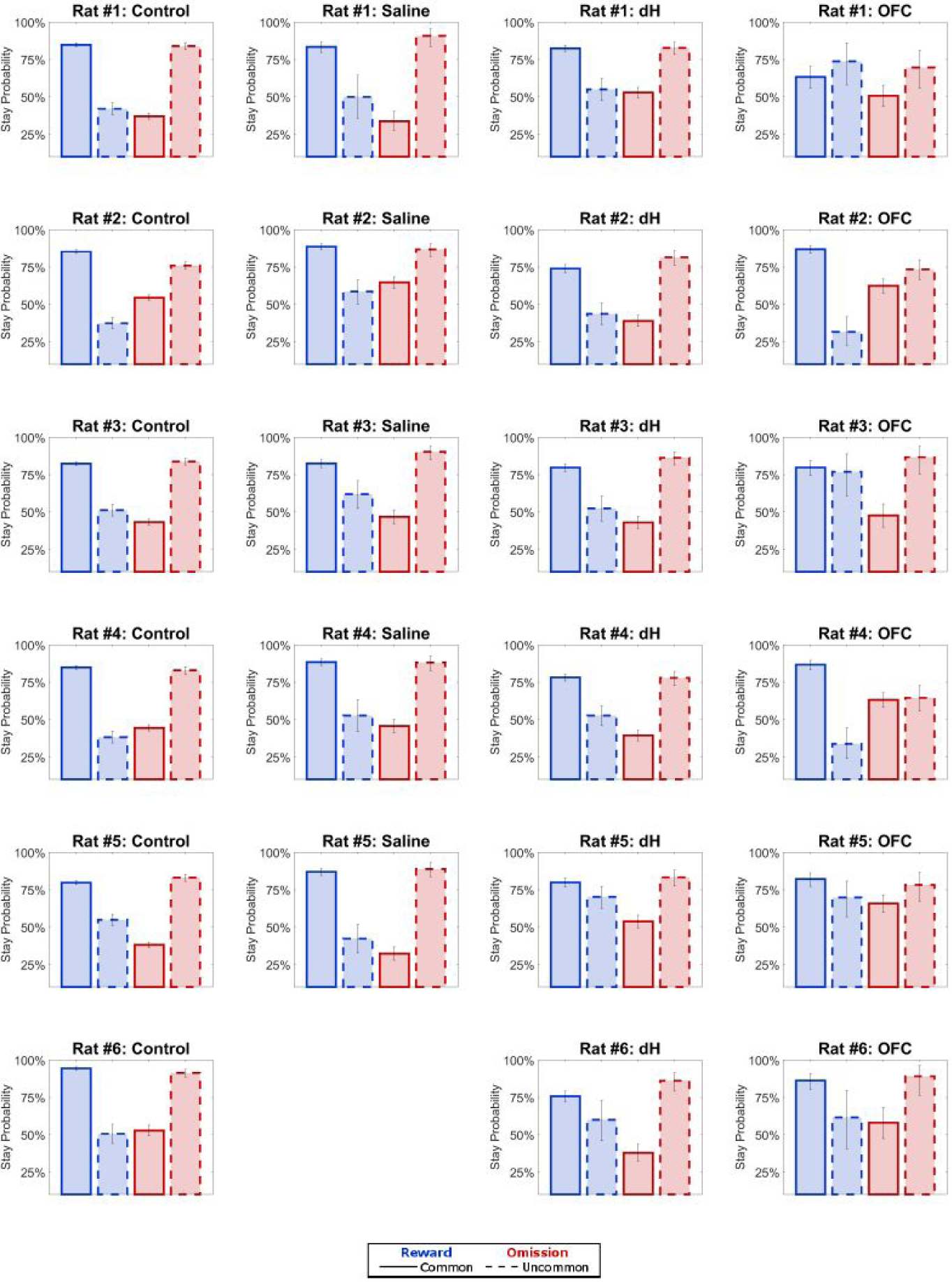
Results of one-trial-back analysis applied to each rat in the inactivation dataset

**Figure S14:**
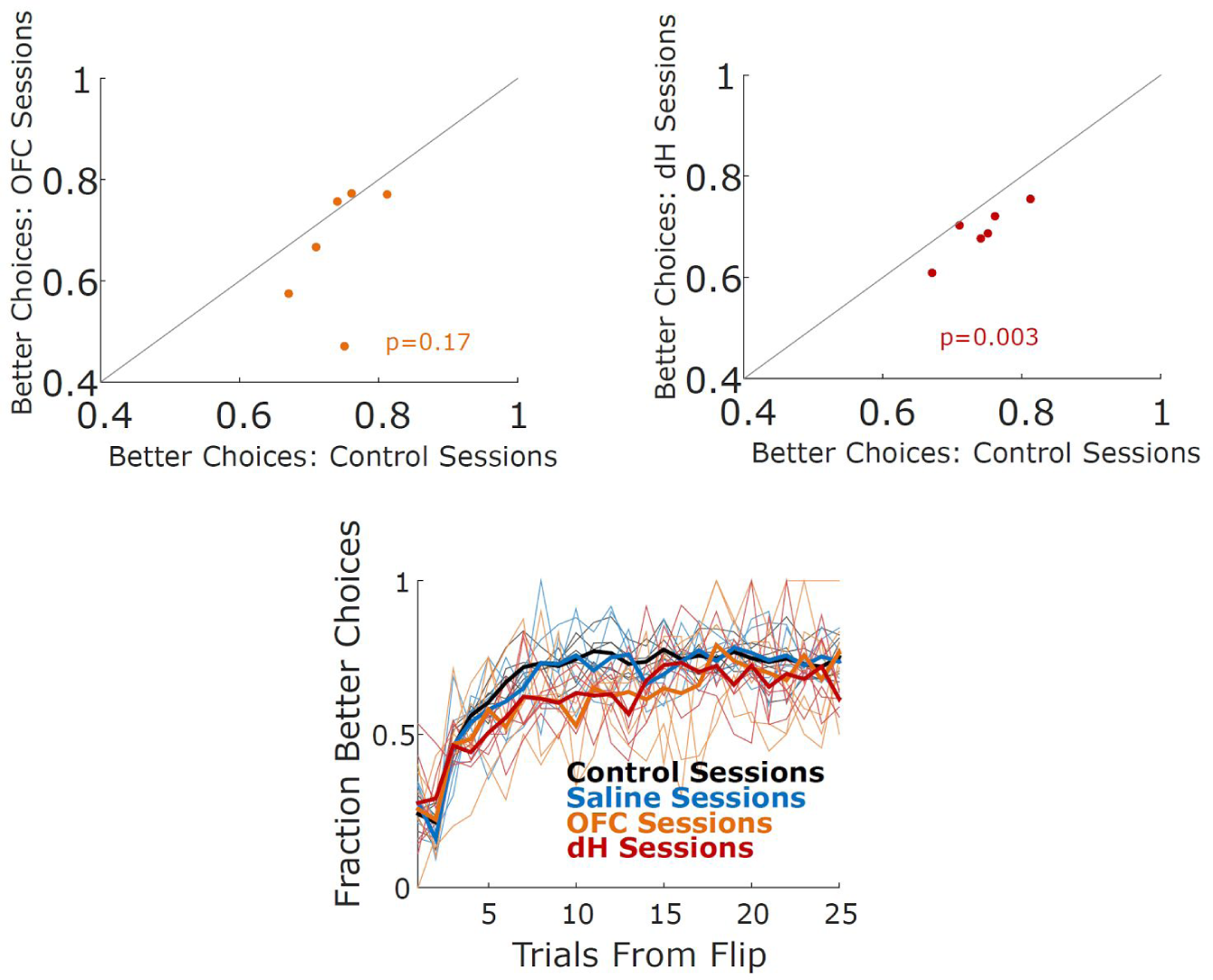
Rat performance compared between inactivation and control sessions. Top: Fraction of times each rat selected the choice port with the greatest probability of leading to the reward port with the greatest probability of reward, for control vs. OFC sessions (Left) and for control vs. dH sessions (Right). Bottom: Fraction of times the better port was selected, as a function of the number of trials since the last reward probability flip.

**Figure S15:**
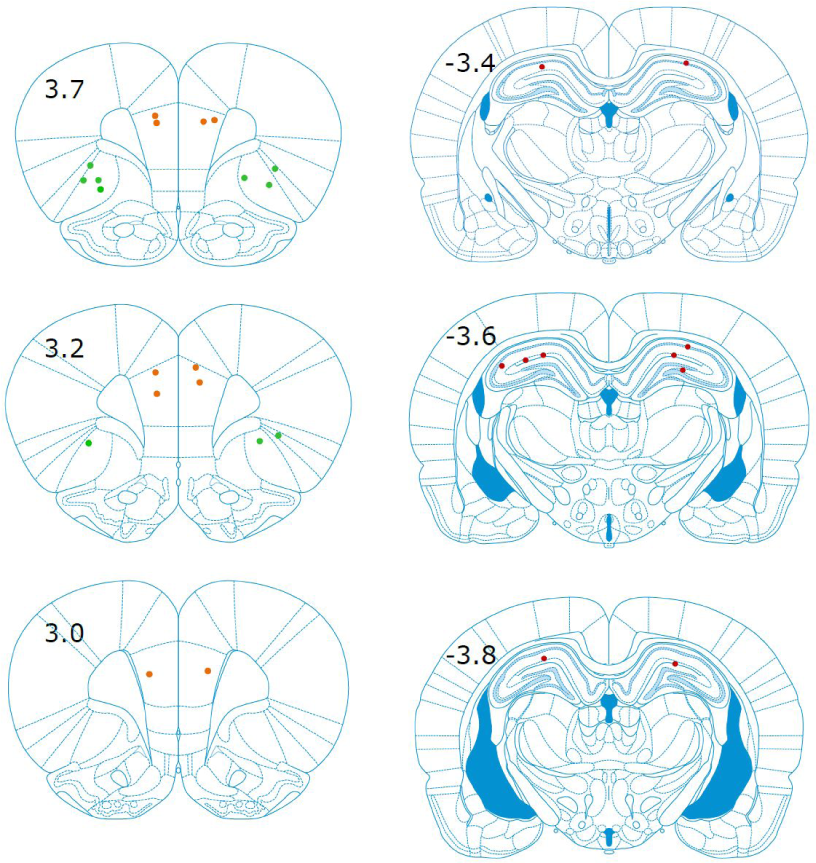
Placement of cannula in individual rats. Green points indicate OFC cannula tips, orange points indicate PL cannula tips, and red points indicate dH cannula tips.

## Supplemental Discussion: Akam, Costa, & Dayan, 2015

Recent work involving extensive analysis of synthetic datasets from various software agents performing the two-step task has suggested that a variety of model-free agents can masquerade as model-based, taking advantage of expanded state space representations (Akam, Costa, & Dayan, 2015). These agents present an important challenge to work using the two-step task to diagnose model-based behavior, since they suggest that planning-typical patterns of choices may not necessarily require the representation of action-outcome relationships. In some cases, this challenge can be fully met using more careful analysis of choice behavior, such as the trial history regression that we employ here. In other cases, it cannot to our knowledge be categorically ruled out, but it can be strongly mitigated by analysis of movement times. In this section, we describe these agents and why we believe that neither they nor any other model-free account provide a likely explanation of our data.

The first of these agents, termed the “reward-as-cue” agent by Akam et al., adopts an expanded representation of state. Where a traditional model-free agent assigns values to the left and right choices, this agent assigns different values to each choice based on the last trial’s outcome. It might learn, for example, a low value for the left choice port following a reward at the left reward port, but a high value following a reward at the right choice port. Over time, this agent can learn to switch its behavior in a similar way as a model-based agent, but only with respect to the outcome of the immediately previous trial. Analysis which considers only this immediately previous trial is therefore unable to distinguish synthetic datasets generated by this agent from those generated by a genuine planning agent. An approach, such as the one we adopt here, which considers the influence of many past trials reveals a key difference between this agent and a planning agent: only the planning agent switches its behavior in a model-based manner with respect to outcomes two or more trials in the past (Miller, Brody, & Botvinick, *bioRxiv*, 2016). Since we see this pattern in our dataset (Figure. 2 and S4), we do not believe that the “reward-as-cue” agent provides a plausible account of behavior on the rat two-step task.

The second agent, termed the “latent state” agent, is aware of the blockwise structure of the task (i.e. that one port is rewarded at 80% and the other at 20%, and that they switch in blocks). The agent performs Bayesian inference after each trial to estimate the likelihood that one block or the other is currently active, and then uses a fixed mapping from its estimate of this likelihood onto choice behavior. It can be described as “model-based” in the sense that this inference requires a model of the dynamics of the world, but not as “planning”, since this model contains no action-outcome information. With an appropriate fixed policy (which might potentially be learned slowly over the course of training by a model-free mechanism), this agent can masquerade as a planning agent even when viewed through the lens of the trial-history regression analysis. In the two-step task, it is in fact a notational variant on a genuine planning agent – the “HMM learner” we consider in Figure 3b – which performs similar latent state inference, but then uses these latent states to plan. This means that no analysis of choice behavior can reveal the difference between synthetic datasets generated by the model-free latent state agent and a genuine planning agent. Key differences, however, are expected in movement times. By definition, only planning agents are aware of the action-outcome contingencies built into the task, and therefore only they have the ability to anticipate the common transition and to respond more quickly on common vs. on uncommon transition trials.

In our dataset, we find a strong effect of outcome probability on movement time – rats show much shorter latency to respond following a common transition than following an uncommon transition (Figure S3). Importantly, in our version of the task, rats must enter the bottom center port before the LEDs cue them which reward port to enter on that trial; the physical movement towards the reward port is therefore the same regardless of which choice port was selected. This limits the possibility that the movement time difference is an artifact of motor facilitation, and argues in favor of the idea that rats are aware of the transition probabilities and anticipate the common transition, using this knowledge to select and prepare actions. Model-free agents, including the latent state agent, by definition are unaware of these contingencies, and would not be expected to show such movement time effects. We therefore believe that any parsimonious account of our data, considered as a whole, will be one in which rats are aware of the action-outcome contingencies built into the task, and in which they use this knowledge both to anticipate transitions and to guide choice behavior – that is, to plan.

We wish to be clear, however, that our data do not categorically rule out two less-parsimonious accounts involving the latent state agent. In the first account, choices are computed by the latent state mechanism, and the movement time effects we observe are due to some type of motor facilitation artifact which is resistant to the constraints of task geometry. In the second account, a latent-state mechanism determines choice at the first step, after which another mechanism (with knowledge of the transition probabilities) guides behavior for the remainder of the trial. Concerns involving the latent state agent apply to all versions of the two-step task, and can most conclusively be resolved by experiments employing neural recordings to investigate neural representations directly. By bringing the two-step task to rodents, we have opened the door to experiments of this kind.

## References

1. Sutton, R. S. & Barto, A. G. Reinforcement learning: An introduction. 1, (MIT press Cambridge, 1998).

2. Tolman, E. C. Cognitive maps in rats and men. Psychol. Rev. 55, 189–208 (1948).

3. Dolan, R. J. & Dayan, P. Goals and habits in the brain. Neuron 80, 312–325 (2013).

4. Balleine, B. W. & O’Doherty, J. P. Human and rodent homologies in action control: corticostriatal determinants of goal-directed and habitual action. Neuropsychopharmacology 35, 48–69 (2010).

5. Daw, N. D., Niv, Y. & Dayan, P. Uncertainty-based competition between prefrontal and dorsolateral striatal systems for behavioral control. Nat. Neurosci. 8, 1704–1711 (2005).

6. Brogden, W. J. Sensory pre-conditioning. J. Exp. Psychol. 25, 323 (1939).

7. Hammond, L. J. The effect of contingency upon the appetitive conditioning of free-operant behavior. J. Exp. Anal. Behav. 34, 297–304 (1980).

8. Adams, C. D. & Dickinson, A. Instrumental responding following reinforcer devaluation. The Quarterly Journal of Experimental Psychology Section B 33, 109–121 (1981).

9. Hilário, M. R. F., Clouse, E., Yin, H. H. & Costa, R. M. Endocannabinoid signaling is critical for habit formation. Front. Integr. Neurosci. 1, 6 (2007).

10. Daw, N. D., Gershman, S. J., Seymour, B., Dayan, P. & Dolan, R. J. Model-based influences on humans’ choices and striatal prediction errors. Neuron 69, 1204–1215 (2011).

11. Simon, D. A. & Daw, N. D. Neural correlates of forward planning in a spatial decision task in humans. J. Neurosci. 31, 5526–5539 (2011).

12. Wunderlich, K., Dayan, P. & Dolan, R. J. Mapping value based planning and extensively trained choice in the human brain. Nat. Neurosci. 15, 786–791 (2012).

13. Lee, S. W., Shimojo, S. & O’Doherty, J. P. Neural computations underlying arbitration between model-based and model-free learning. Neuron 81, 687–699 (2014).

14. O’keefe, J. & Nadel, L. The hippocampus as a cognitive map. 3, (Clarendon Press Oxford, 1978).

15. Packard, M. G. & McGaugh, J. L. Inactivation of hippocampus or caudate nucleus with lidocaine differentially affects expression of place and response learning. Neurobiol. Learn. Mem. 65, 65–72 (1996).

16. Morris, R. G., Garrud, P., Rawlins, J. N. & O’Keefe, J. Place navigation impaired in rats with hippocampal lesions. Nature 297, 681–683 (1982).

17. O’Keefe, J. & Dostrovsky, J. The hippocampus as a spatial map. Preliminary evidence from unit activity in the freely-moving rat. Brain Res. 34, 171–175 (1971).

18. Johnson, A. & Redish, A. D. Neural ensembles in CA3 transiently encode paths forward of the animal at a decision point. J. Neurosci. 27, 12176–12189 (2007).

19. Wikenheiser, A. M. & Redish, A. D. Hippocampal theta sequences reflect current goals. Nat. Neurosci. 18, 289–294 (2015).

20. Diba, K. & Buzsáki, G. Forward and reverse hippocampal place-cell sequences during ripples. Nat. Neurosci. 10, 1241–1242 (2007).

21. Pfeiffer, B. E. & Foster, D. J. Hippocampal place-cell sequences depict future paths to remembered goals. Nature 497, 74–79 (2013).

22. Koene, R. A., Gorchetchnikov, A., Cannon, R. C. & Hasselmo, M. E. Modeling goal-directed spatial navigation in the rat based on physiological data from the hippocampal formation. Neural Netw. 16, 577–584 (2003).

23. Johnson, A., van der Meer, M. A. A. & Redish, A. D. Integrating hippocampus and striatum in decision-making. Curr. Opin. Neurobiol. 17, 692–697 (2007).

24. Foster, D. J. & Knierim, J. J. Sequence learning and the role of the hippocampus in rodent navigation. Curr. Opin. Neurobiol. 22, 294–300 (2012).

25. Pezzulo, G., van der Meer, M. A. A., Lansink, C. S. & Pennartz, C. M. A. Internally generated sequences in learning and executing goal-directed behavior. Trends Cogn. Sci. 18, 647–657 (2014).

26. Kimble, D. P. & BreMiller, R. Latent learning in hippocampal-lesioned rats. Physiol. Behav. 26, 1055–1059 (1981).

27. Kimble, D. P., Jordan, W. P. & BreMiller, R. Further evidence for latent learning in hippocampal-lesioned rats. Physiol. Behav. 29, 401–407 (1982).

28. Corbit, L. H. & Balleine, B. W. The role of the hippocampus in instrumental conditioning. J. Neurosci. 20, 4233–4239 (2000).

29. Corbit, L. H., Ostlund, S. B. & Balleine, B. W. Sensitivity to instrumental contingency degradation is mediated by the entorhinal cortex and its efferents via the dorsal hippocampus. J. Neurosci. 22, 10976–10984 (2002).

30. Ward-Robinson, J. et al. Excitotoxic lesions of the hippocampus leave sensory preconditioning intact: Implications for models of hippocampal functioning. Behav. Neurosci. 115, 1357 (2001).

31. Gaskin, S., Chai, S.-C. & White, N. M. Inactivation of the dorsal hippocampus does not affect learning during exploration of a novel environment. Hippocampus 15, 1085–1093 (2005).

32. Bunsey, M. & Eichenbaum, H. Conservation of hippocampal memory function in rats and humans. Nature 379, 255–257 (1996).

33. Dusek, J. A. & Eichenbaum, H. The hippocampus and memory for orderly stimulus relations. Proc. Natl. Acad. Sci. U. S. A. 94, 7109–7114 (1997).

34. Van der Jeugd, A. et al. Hippocampal involvement in the acquisition of relational associations, but not in the expression of a transitive inference task in mice. Behav. Neurosci. 123, 109–114 (2009).

35. Devito, L. M. & Eichenbaum, H. Memory for the order of events in specific sequences: contributions of the hippocampus and medial prefrontal cortex. J. Neurosci. 31, 3169–3175 (2011).

36. Jones, J. L. et al. Orbitofrontal cortex supports behavior and learning using inferred but not cached values. Science 338, 953–956 (2012).

37. Gallagher, M., McMahan, R. W. & Schoenbaum, G. Orbitofrontal cortex and representation of incentive value in associative learning. J. Neurosci. 19, 6610–6614 (1999).

38. McDannald, M. A., Lucantonio, F., Burke, K. A., Niv, Y. & Schoenbaum, G. Ventral striatum and orbitofrontal cortex are both required for model-based, but not model-free, reinforcement learning. J. Neurosci. 31, 2700–2705 (2011).

39. Gremel, C. M. & Costa, R. M. Orbitofrontal and striatal circuits dynamically encode the shift between goal-directed and habitual actions. Nat. Commun. 4, 2264 (2013).

40. Miller, K. J., Brody, C. D. & Botvinick, M. M. Identifying Model-Based and Model-Free Patterns in Behavior on Multi-Step Tasks. bioRxiv 096339 (2016). doi:10.1101/096339

41. Krupa, D. J., Ghazanfar, A. A. & Nicolelis, M. A. Immediate thalamic sensory plasticity depends on corticothalamic feedback. Proc. Natl. Acad. Sci. U. S. A. 96, 8200–8205 (1999).

42. Martin, J. H. Autoradiographic estimation of the extent of reversible inactivation produced by microinjection of lidocaine and muscimol in the rat. Neurosci. Lett. 127, 160–164 (1991).

43. Aarts, E., Verhage, M., Veenvliet, J. V., Dolan, C. V. & van der Sluis, S. A solution to dependency: using multilevel analysis to accommodate nested data. Nat. Neurosci. 17, 491–496 (2014).

44. Gelman, A. et al. Bayesian Data Analysis, Third Edition. (CRC Press, 2013).

45. Daw, N. D. in Decision Making, Affect, and Learning 3–38 (2011).

46. Akam, T., Costa, R. & Dayan, P. Simple Plans or Sophisticated Habits? State, Transition and Learning Interactions in the Two-Step Task. PLoS Comput. Biol. 11, e1004648 (2015).

47. Smittenaar, P., FitzGerald, T. H. B., Romei, V., Wright, N. D. & Dolan, R. J. Disruption of dorsolateral prefrontal cortex decreases model-based in favor of model-free control in humans. Neuron 80, 914–919 (2013).

48. Otto, A. R., Gershman, S. J., Markman, A. B. & Daw, N. D. The Curse of Planning: Dissecting Multiple Reinforcement-Learning Systems by Taxing the Central Executive. Psychol. Sci. 24, 751–761 (2013).

49. Wunderlich, K., Smittenaar, P. & Dolan, R. J. Dopamine enhances model-based over model-free choice behavior. Neuron 75, 418–424 (2012).

50. Economides, M., Kurth-Nelson, Z., Lübbert, A., Guitart-Masip, M. & Dolan, R. J. Model-Based Reasoning in Humans Becomes Automatic with Training. PLoS Comput. Biol. 11, e1004463 (2015).

51. Keramati, M., Dezfouli, A. & Piray, P. Speed/accuracy trade-off between the habitual and the goal-directed processes. PLoS Comput. Biol. 7, e1002055 (2011).

52. Kool, W., Cushman, F. A. & Gershman, S. J. When does model-based control pay off? PLOS Computational Biology (2016).

53. Izquierdo, A., Suda, R. K. & Murray, E. A. Bilateral orbital prefrontal cortex lesions in rhesus monkeys disrupt choices guided by both reward value and reward contingency. J. Neurosci. 24, 7540–7548 (2004).

54. Machado, C. J. & Bachevalier, J. The effects of selective amygdala, orbital frontal cortex or hippocampal formation lesions on reward assessment in nonhuman primates. Eur. J. Neurosci. 25, 2885–2904 (2007).

55. Padoa-Schioppa, C. Neurobiology of economic choice: a good-based model. Annu. Rev. Neurosci. 34, 333–359 (2011).

56. Schoenbaum, G., Roesch, M. R., Stalnaker, T. A. & Takahashi, Y. K. A new perspective on the role of the orbitofrontal cortex in adaptive behaviour. Nat. Rev. Neurosci. 10, 885–892 (2009).

57. Wilson, R. C., Takahashi, Y. K., Schoenbaum, G. & Niv, Y. Orbitofrontal cortex as a cognitive map of task space. Neuron 81, 267–279 (2014).

58. Ostlund, S. B. & Balleine, B. W. Orbitofrontal cortex mediates outcome encoding in Pavlovian but not instrumental conditioning. J. Neurosci. 27, 4819–4825 (2007).

59. Foster, D. J., Morris, R. G. & Dayan, P. A model of hippocampally dependent navigation, using the temporal difference learning rule. Hippocampus 10, 1–16 (2000).

60. Olton, D. S., Becker, J. T. & Handelmann, G. E. Hippocampus, space, and memory. Behav. Brain Sci. 2, 313–322 (1979).

61. Racine, R. J. & Kimble, D. P. Hippocampal lesions and delayed alternation in the rat. Psychon. Sci. 3, 285–286 (1965).

62. Olton, D. S. & Papas, B. C. Spatial memory and hippocampal function. Neuropsychologia 17, 669–682 (1979).

63. Olton, D. S., Walker, J. A. & Gage, F. H. Hippocampal connections and spatial discrimination. Brain Res. 139, 295–308 (1978).

64. Gilboa, A., Sekeres, M., Moscovitch, M. & Winocur, G. Higher-order conditioning is impaired by hippocampal lesions. Curr. Biol. 24, 2202–2207 (2014).

65. McEchron, M. D., Bouwmeester, H., Tseng, W., Weiss, C. & Disterhoft, J. F. Hippocampectomy disrupts auditory trace fear conditioning and contextual fear conditioning in the rat. Hippocampus 8, 638–646 (1998).

66. Solomon, P. R., Vander Schaaf, E. R., Thompson, R. F. & Weisz, D. J. Hippocampus and trace conditioning of the rabbit’s classically conditioned nictitating membrane response. Behav. Neurosci. 100, 729–744 (1986).

67. O’Keefe, J. Do hippocampal pyramidal cells signal non-spatial as well as spatial information? Hippocampus 9, 352–364 (1999).

68. Burgess, N. The 2014 Nobel Prize in Physiology or Medicine: a spatial model for cognitive neuroscience. Neuron 84, 1120–1125 (2014).

69. Hartley, T., Lever, C., Burgess, N. & O’Keefe, J. Space in the brain: how the hippocampal formation supports spatial cognition. Philos. Trans. R. Soc. Lond. B Biol. Sci. 369, 20120510 (2014).

70. Eichenbaum, H., Dudchenko, P., Wood, E., Shapiro, M. & Tanila, H. The hippocampus, memory, and place cells: is it spatial memory or a memory space? Neuron 23, 209–226 (1999).

71. Buckner, R. L. The role of the hippocampus in prediction and imagination. Annu. Rev. Psychol. 61, 27–48, C1–8 (2010).

72. Eichenbaum, H. & Cohen, N. J. Can we reconcile the declarative memory and spatial navigation views on hippocampal function? Neuron 83, 764–770 (2014).

73. Schiller, D. et al. Memory and Space: Towards an Understanding of the Cognitive Map. J. Neurosci. 35, 13904–13911 (2015).

74. Lau, B. & Glimcher, P. W. Dynamic response-by-response models of matching behavior in rhesus monkeys. J. Exp. Anal. Behav. 84, 555–579 (2005).

75. Stan Development Team. MatlabStan: The MATLAB interface to Stan. (2016).

76. Bob Carpenter, Andrew Gelman, Matt Hoffman, Daniel Lee, Ben Goodrich, Michael Betancourt, Michael A. Brubaker, Jiqiang Guo, Peter Li, and Allen Riddell. Stan: A Probabilistic Programming Language. (2016).

77. Duane, S., Kennedy, A. D., Pendleton, B. J. & Roweth, D. Hybrid Monte Carlo. Phys. Lett. B 195, 216–222 (1987).

